# Neural signatures of attentional engagement during narratives and its consequences for event memory

**DOI:** 10.1101/2020.08.26.266320

**Authors:** Hayoung Song, Emily S. Finn, Monica D. Rosenberg

## Abstract

As we comprehend narratives, our attentional engagement fluctuates over time. Despite theoretical conceptions of narrative engagement as emotion-laden attention, little empirical work has characterized the cognitive and neural processes that comprise subjective engagement in naturalistic contexts or its consequences for memory. Here, we relate fluctuations in narrative engagement to patterns of brain coactivation, and test whether neural signatures of engagement predict later recall. In behavioral studies, participants continuously rated how engaged they were as they watched a television episode or listened to a story. Self-reported engagement was synchronized across individuals and driven by the emotional content of the narratives. During fMRI, we observed highly synchronized activity in the default mode network when people were, on average, more engaged in the same narratives. Models based on time-varying whole-brain functional connectivity predicted evolving states of engagement across participants and even across different datasets. The same functional connections also predicted post-scan event recall, suggesting that engagement during encoding impacts subsequent memory. Finally, group-average engagement was related to fluctuations of an independent functional connectivity index of sustained attention. Together, our findings characterize the neural signatures of engagement dynamics and elucidate relationships between narrative engagement, sustained attention, and event memory.

## Introduction

We engage with the world and construct memories by attending to information in our external environment.^1^ However, the degree to which we pay attention waxes and wanes over time.^2,3^ Such fluctuations of attention not only influence our ongoing perceptual experience, but can also have consequences for what we later remember.^4^

Changes in attentional states are typically studied with continuous performance tasks (CPTs), which require participants to respond to rare targets in a constant stream of stimuli or respond to every presented stimulus *except* the rare target.^5–7^ Paying attention to taxing CPTs, however, often *feels* different than paying attention in other everyday situations. For example, when we listen to the radio, watch a television show, or have a conversation with family and friends, sustaining focus can feel comparatively effortless. Psychology research has characterized feelings of effortless attention in other contexts, such as *flow* states of complete absorption in an activity.^8^ When comprehending narratives, our attention may be naturally captured by the story, causing us to become engaged in the experience. Narrative engagement has been defined as an experience of being deeply immersed in a story with heightened emotional arousal and attentional focus.^9,10^ Building on this theoretical definition, we characterize how subjective engagement fluctuates as narratives unfold, and test the hypothesis that engagement scales with a story’s emotional content as well as an individual’s sustained attentional state.

Functional neuroimaging studies have used naturalistic, narrative stimuli to examine how we perceive^11,12^ and remember^13^ structured events based on memory of contexts,^14–16^ prior knowledge or beliefs,^17–19^ and emotional and social reasoning.^20–22^ However, strikingly few neuroimaging studies have directly probed attention during naturalistic paradigms. Among these, Regev et al.^23^ examined how selectively attending to narrative inputs from a particular sensory modality (e.g., auditory) while suppressing the other modality (e.g., visual) enhances stimulus-locked brain responses to the attended inputs. With non-narrative movies, Çukur et al.^24^ found that semantic representations were warped towards attended object categories during visual search. However, both of these studies relied on experimental manipulations of attention to elucidate its relationship to ongoing brain activity. Recent studies hypothesized the role of attention dynamics in story comprehension, but did not empirically measure attentional states.^25,26^ Dmochowski et al.^27,28^ showed that inter-subject electroencephalogram (EEG) synchrony (i.e., neural reliability) increases during arousing moments of narrative films, and that indicators of population-level engagement can be predicted from neural reliability. However, the ways in which attention changes in narrative contexts—and the consequences that this has for story comprehension and memory—remain poorly understood.

Here we characterize attentional states in real-world settings by tracking subjective engagement during movie watching and story listening. In doing so, we address three primary aims: testing the theoretical conception of engagement as emotion-laden attention, examining how engagement is reflected in brain dynamics, and elucidating the consequences of engagement during encoding for subsequent memory. We first measured self-reported engagement as behavioral participants watched an episode of television series *Sherlock* or listened to an audio-narrated story, *Paranoia*. Providing empirical support for its theoretical definition, changes in engagement were driven by the emotional contents of narratives and related to fluctuations of a validated functional connectivity (FC) index of sustained attention during psychological tasks. We next related group-average engagement time-courses to fMRI activity observed as a separate pool of participants watched *Sherlock*^13^ or listened to *Paranoia*.^18^ Dynamic inter-subject correlation analysis (ISC)^29^ revealed that activity in large-scale functional networks, especially the default mode network (DMN) was more synchronized across individuals during more engaging periods of the narratives. Further, using time-resolved predictive modeling,^25^ we found that patterns of time-resolved FC predicted engagement dynamics, and that these same patterns predicted later event recall. Thus, we provide evidence for engagement as emotion-laden sustained attention, elucidate the role of brain network dynamics in engagement, and demonstrate relationships between engagement and long-term episodic memory.

## Results

### Tracking dynamic states of engagement during movie watching and story listening

We asked how subjective engagement changes over time as individuals comprehend a story, and whether changes are synchronized across participants. We measured engagement during two narrative stimuli that were used in previous fMRI research: i) a 20m audio-narrated story, *Paranoia* (fMRI dataset *n* = 22),^18^ which was intentionally created to induce suspicion surrounding the characters and situations, and ii) a 50m episode of BBC’s television series *Sherlock* (fMRI dataset *n* = 17),^13^ in which Sherlock and Dr. Watson meet and solve a mysterious crime together. We chose these stimuli because they were both long narratives with complex plots and relationships among characters, and they involved different sensory modalities (visual and auditory), which enabled us to investigate high-level cognitive states of engagement that are independent of sensory modality.

In behavioral studies, as participants listened to *Paranoia* (*n* = 21) or watched *Sherlock* (*n* = 17), they were instructed to continuously rate how engaging they found the story by adjusting a scale bar from 1 (“Not engaging at all”) to 9 (“Completely engaging”) using button presses (**Fig. 1a**). The definition of engagement (**Table 1**; adapted from Matthews et al.^30^) was explained to participants prior to the task. The scale bar was always visible on the computer monitor so that participants could make continuous adjustments as they experienced changes in subjective engagement.

**Figure 1.**
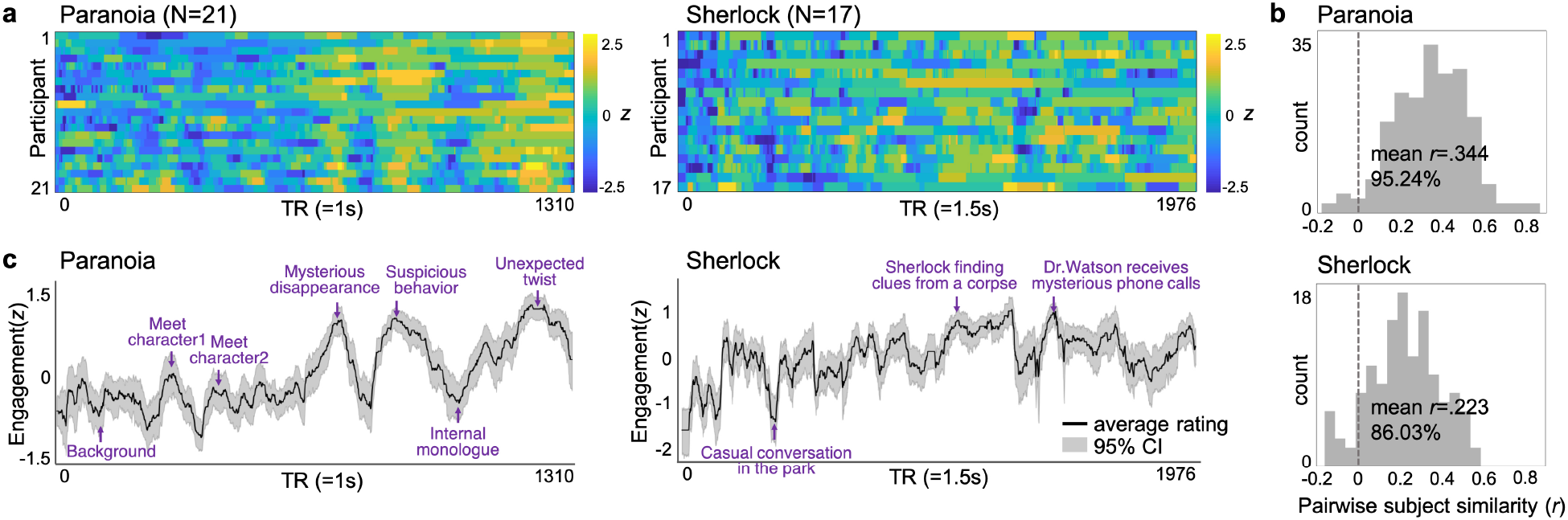
Behavioral experiment results. Ratings of subjective engagement collected as participants listened to the *Paranoia* story (left) or watched the *Sherlock* episode (right). (**a**) Every participant’s engagement ratings across time. Ratings were z-normalized across time for each participant. (**b**) Histograms of pairwise participants’ response similarities. Pearson’s correlations were Fisher’s *r*-to-*z* transformed, and the significance of the correlations was corrected for multiple comparisons using FDR correction. (**c**) Averaged engagement ratings, which are used as proxies for group-level states of engagement. The gray area indicates 95% confidence interval (CI). Event descriptions at the moments of peak engagement are written in purple, with the time indices indicated with arrows.

**Table 1.**
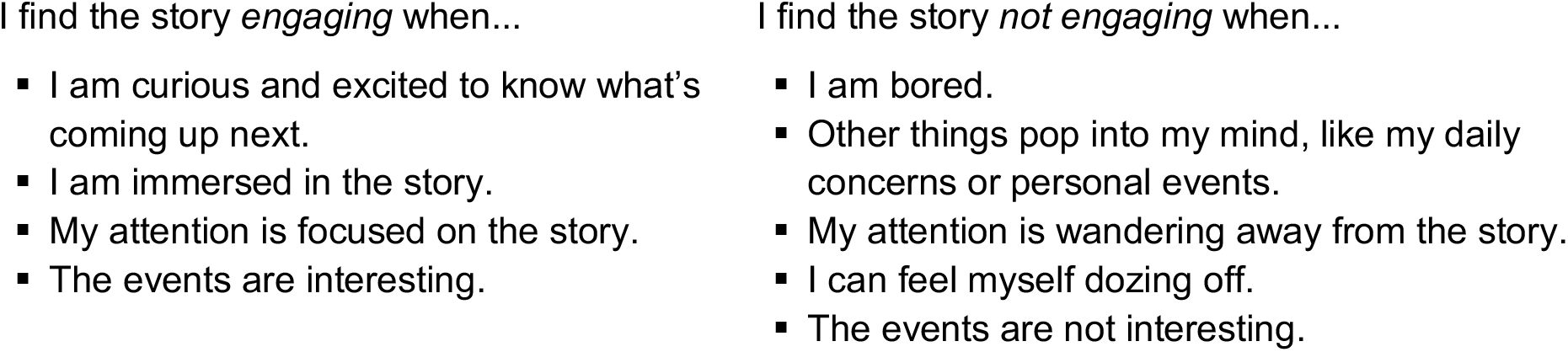
Definition of engagement instructed to the participants during the behavioral experiments.

We first tested whether changes in engagement were similar across participants comprehending the same stories. Providing initial validation for our self-report task as a measure of stimulus-related narrative engagement, we found significant positive correlations between pairwise engagement time-courses (*Paranoia* mean Pearson’s *r* = .344 ± 0.179; *Sherlock* mean *r* = .223 ± 0.170; **Fig. 1b**). 95.24% of *Paranoia* participant pairs and 86.03% of *Sherlock* participant pairs exhibited significant positive correlations (FDR-corrected *p* < .05 after Fisher’s *r*-to-*z* transformation). We averaged all participants’ response time-courses (**Fig. 1c**) and observed positive linear trends in engagement for both *Paranoia* (*t*(1308) = 40.62,*r*^2^= 0.56, *p* < .001) and *Sherlock* (*t*(1974) = 26.36, *r*^2^= 0.26, *p* < .001), suggesting a gradual increase in engagement as the narrative develops to an end.

Next, to assess the consistency of engagement ratings during different parts of the narratives, we calculated pairwise participants’ response consistency for each of the three runs of *Paranoia* and two runs of *Sherlock*. The stimuli were segmented with interim breaks so that the procedure matched experimental runs of the previously collected fMRI studies. Consistency of engagement ratings increased over time for *Paranoia* (run 1 mean Pearson’s *r* = .05 ± .27, run 2 = .38 ± .24, run 3 = .41 ± .30; repeated measures ANOVA: F(2,418) = 121.26, *p* < .001) but decreased over time for *Sherlock* (run 1 = .22 ± .22, run 2 = .11 ± .28; F(1,135) = 15.25, *p* < .001), suggesting that the degree to which engagement is shared across individuals is idiosyncratic to different stories and may depend on narrative content.

Since engagement ratings were shared across participants, we treat group-average engagement time-courses (**Fig. 1c**) as a proxy for stimulus-related engagement, common across individuals. We qualitatively assessed the moments when participants were, on average, most engaged or less engaged in the narratives (**Fig. 1c**). In *Paranoia*, engagement peaked at moments when a character exhibited suspicious behavior, or when there was an unexpected twist in the story. On the other hand, engagement was generally low when a story setting was being developed or when a protagonist was having an internal thought. In *Sherlock*, engagement peaked at moments when Sherlock was solving a mysterious crime and when events were highly suspenseful. Participants’ general engagement decreased during comparatively casual events, less relevant to the crime scenes. The averaged engagement time-courses were convolved with the hemodynamic response function (HRF) to be applied to a separate pool of individuals who participated in previous fMRI studies.

### Engagement scales with emotional arousal of the narratives

Given that participants’ engagement dynamics were time-locked to the stimuli, we asked which features of narratives drove changes in engagement. We conducted partial correlations between group-average engagement and four features of the narratives: positive and negative emotional content, and auditory and visual sensory information measured at every moment of time. We used different ways of extracting the emotional content of the *Paranoia* and *Sherlock* narratives to utilize open resources distributed by the two research groups and assess the robustness of results to specific extraction approach. For *Paranoia*, emotional arousal was inferred from the occurrence of positive and negative emotion words in the story transcript using Linguistic Inquiry and Word Count (LIWC) text analysis^31^ (provided by Finn et al.^18^). Words indicating positive emotion included *accept*, *adventur*e*, amazingly*, and *appreciated*, and words indicating negative emotion included *afraid*, *alarmed*, *anxiou*s, and *apprehensively* (for a full list of emotional words, see **Supplementary Table 1**). The frequencies of positive and negative emotion words were counted per sentence, then expanded to match the moments when the sentence was uttered during scans (**Supplementary Fig. 1a**). Thus, high frequencies indicated high positive or negative emotional arousal. For *Sherlock*, we used four independent raters’ emotional ratings (provided by Chen et al.^13^) at each scene of *Sherlock*, containing both valence (positive and negative) and arousal (scale of 1 to 5). Inter-rater reliability was high, with average of the pairwise raters’ Pearson’s *r* = 0.542 ± 0.059, thus we used an averaged rating as a proxy for evolving emotional arousal, again separately for positive and negative emotional content. To quantify auditory salience of the stimulus, we extracted audio envelopes from the sound clips of both *Paranoia* and *Sherlock* using Hilbert transform.^32^ For *Sherlock,* the visual salience was represented by global luminance, i.e., the average of pixel-wise luminance values per frame of the video.^33^

The occurrence of negative, but not positive, emotion words in *Paranoia* was significantly correlated with changes in engagement (positive: partial *r* = −.019, *p* = .859; negative: partial *r* = .133, *p* = .018) when controlling for the rest of the variables (**Supplementary Fig. 1b**). In *Sherlock*, both positive (partial *r* = .530, *p* < .001) and negative (partial *r* = .539, *p* < .001) emotional arousal were correlated with engagement. We observed no significant relationship between auditory envelopes and engagement, both for *Paranoia* (partial *r* = −.018, *p* = .451) and *Sherlock* (partial *r* = −.103, *p* = .373). Similarly, there was no significant correlation between visual luminance and engagement for *Sherlock* (partial *r* = −.230, *p* = .096). For significance tests, observed partial correlation values were compared with null distributions of partial correlations between each factor and 10,000 permuted, phase randomized engagement ratings (two-tailed test: *p* = (1 + number of null |*r*| values ≥ empirical |*r*|) / (1 + number of permutations)). We created such null distribution because phase randomization retains the same characteristics of temporal dynamics (i.e., frequency and amplitude) but in different phases. The results suggest that engagement scales with emotional narrative content, but not with the sensory-level salience of the stimuli themselves.

### Stimulus-evoked BOLD responses increase during states of engagement

We analyzed open-sourced fMRI data from two separate studies.^13,18^ The datasets were collected from different participant samples, experimental sites, and research contexts, with different image acquisition protocols. To assess the reproducibility and generalizability of our results, we applied different analysis pipelines to the two datasets throughout the study. This allowed us to conceptually replicate findings across samples and confirm that results are robust to particular analytic choices. The *Paranoia* fMRI data were preprocessed with our built-in preprocessing pipeline (see details in *Methods*), whereas we used the fully preprocessed images of *Sherlock* dataset, provided by Chen et al.^13^

We asked whether stimulus-evoked BOLD responses scaled with changes in engagement. Inter-subject correlation (ISC) is a method of isolating shared, stimulus-driven brain activity, assuming that if participants perceive the same stimulus at the same time, their shared variance in fMRI signal is related to stimulus processing.^29,34–36^ Recent studies showed that ISC provides an alternative to identifying brain regions that are entrained to a stimulus with reduced intrinsic noise^35^ compared to a conventional general linear model (GLM) that relies on fixed experimental manipulations with *a priori* hypotheses^29^ (but see **Supplementary Figure 2** for GLM results with continuous group-average engagement as a regressor). We hypothesized that ISC would increase as participants, on average, become more engaged in the story.^28^ Again, we used group-average ratings from the behavioral studies as our index of narrative engagement because measures of engagement were not available in the existing fMRI datasets.

To test the hypothesis that ISC varies with engagement, we calculated dynamic ISC using a tapered sliding window approach, where the Pearson’s correlation between pairwise participants’ BOLD responses was computed within the temporal window, repeatedly across the entire scan duration.^37^ We implemented a window size of TR = 40 (= 40s) for *Paranoia* and TR = 30 (= 45s) for *Sherlock* datasets, following the optimal window size suggested by previous literature,^38–40^ with a step size of 1 TR and a Gaussian kernel σ = 3 TR. The BOLD time-course was extracted from each region of interest (ROI) in Yeo et al.’s^41,42^ 122-ROI parcellation. We selected this atlas because it includes both cortical and subcortical areas of the brain and annotates each ROI with one of the eight functional networks that represent the distributed functional organization of the brain. Dynamic ISC was calculated for all pairs of participants, and was averaged per ROI. The Pearson’s *r* between the dynamic ISC time-course and group-average engagement (smoothed with the same sliding window approach) was calculated for each ROI (**Fig. 2a**).

**Figure 2.**
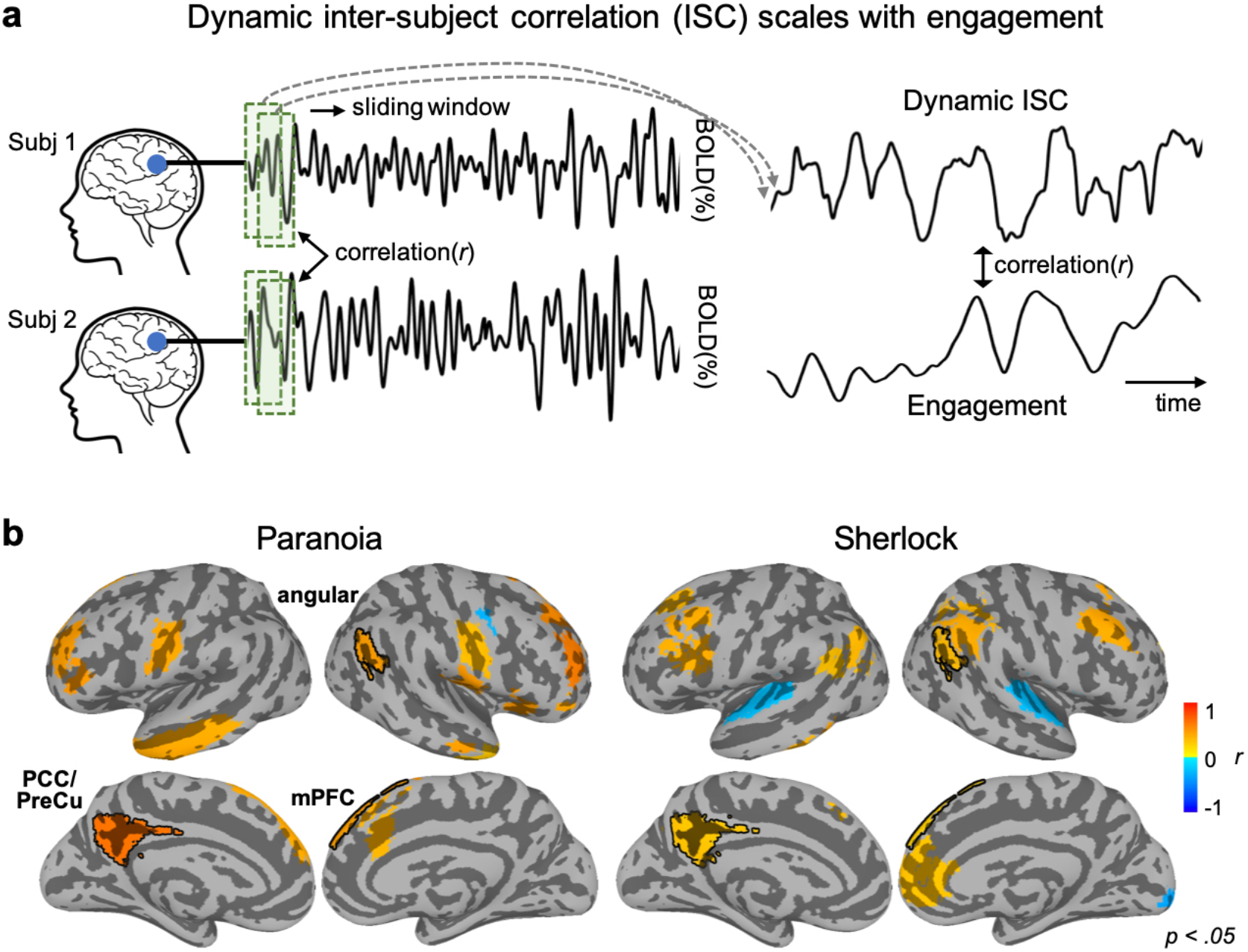
Dynamic inter-subject correlation (ISC) scales with changes in the states of engagement. (**a**) Schematic of dynamic ISC analysis. ISC was calculated by correlating pairwise participants’ ROI time-courses within a time window. Sliding window analysis was used to measure the dynamic changes in the ISC. The dynamic ISCs of all pairwise participants were averaged per region-of-interest (ROI). The correlation between dynamic ISC and group-average engagement was calculated. (**b**) Regions that show significant correlation between dynamic ISC and group-average engagement (uncorrected *p* < 0.05), for *Paranoia* (left) and *Sherlock* (right) datasets. Significance tests were conducted per ROI by comparing the observed correlation value with a permuted distribution in which the engagement rating was phase randomized. Three regions selected in both datasets are indicated with black contours. Angular: angular gyrus, mPFC: medial prefrontal cortex, PCC/PreCu: posterior cingulate cortex/precuneus.

**Fig. 2b** shows the regions in which dynamic ISC significantly correlated with engagement ratings (two-tailed test non-parametric *p* < .05, uncorrected for multiple comparisons). Dynamic ISC in 21 of the 122 Yeo atlas ROIs was significantly correlated with engagement for *Paranoia*, and dynamic ISC in 20 ROIs was significantly correlated with engagement for *Sherlock* (for a full list of significant regions, see **Supplementary Table 2**). In almost all of these regions, dynamic ISC was positively correlated with engagement (20/21 and 17/20 regions, respectively), suggesting that the stimulus-evoked BOLD responses increased as narrative engagement increased. Regions in the default mode network (DMN) were selected above chance for both datasets (8/21 regions, *p* = .042 from non-parametric permutation tests with randomly selected regions, and 9/20 regions corresponded to DMN, *p* = .008, in *Paranoia* and *Sherlock*, respectively), whereas no other Yeo et al.^42^ networks were consistently selected above chance for the two datasets. Furthermore, three regions—all in the DMN—showed higher synchrony with increasing narrative engagement in both the *Paranoia* and *Sherlock* samples: left posterior cingulate cortex [+5.8, +51.0, +31.0] (*r* = .651, *r* = .307 for *Paranoia* and *Sherlock* respectively), right angular gyrus [−51.6, +57.1, +29.1] (*r* = .453, *r* = .316), and right superior medial gyrus [−10.2, −48.4, +41.8] (*r* = .452, *r* = .237). Though the probability of overlap itself was not significant (*p* = .534), the probability that all overlapping regions corresponded to the DMN was above chance (*p* = .031). The results replicated when we used a different parcellation scheme, the Shen atlas (**Supplementary Fig. 3**).^43^ These results suggest that stimulus-evoked BOLD responses increase when subjective levels of engagement increase, especially the regions of the DMN.

### Dynamic prediction of engagement from multivariate patterns of brain activity

Recent work suggests that diverse cognitive and attentional states can be predicted from multivariate patterns of fMRI activity.^44–46^ Thus, we asked whether evolving states of engagement can be predicted from dynamic patterns of brain activity. To this end, we applied dynamic predictive modeling^25^ to BOLD time-courses of ROIs to predict group-average engagement at every moment of time.

Decoding was conducted using a leave-one-subject-out (LOO) cross-validation within each dataset. Support vector regression (SVR) models with an assumption of non-linear feature space were trained using fMRI data from all but one participant, and applied to the held-out participants’ pattern of BOLD activity at every TR to predict the group-average engagement observed at the corresponding TR (**Fig. 3a**). Prediction accuracy was calculated by averaging the Pearson’s correlations between the predicted and observed engagement dynamics across cross-validation folds. We used correlation as an indicator of predictive performance because we were interested in whether the model captures temporal dynamics, rather than the actual values of group-average engagement ratings. Nevertheless, we report both the RMSE and *R*^2^ along with the correlation values. For significance tests, observed prediction accuracy was compared with results from 1,000 permutations where null models were trained and tested with actual brain patterns to predict the phase randomized engagement ratings. We assumed a one-tailed significance test, with *p* = (1 + number of null *r* values ≥ empirical *r*) / (1 + number of permutations) for Pearson’s correlation and *R*^2^, but applied an opposite end of tail for RMSE (i.e., the smaller the RMSE, the better the prediction compared to null hypothesis).

**Figure 3.**
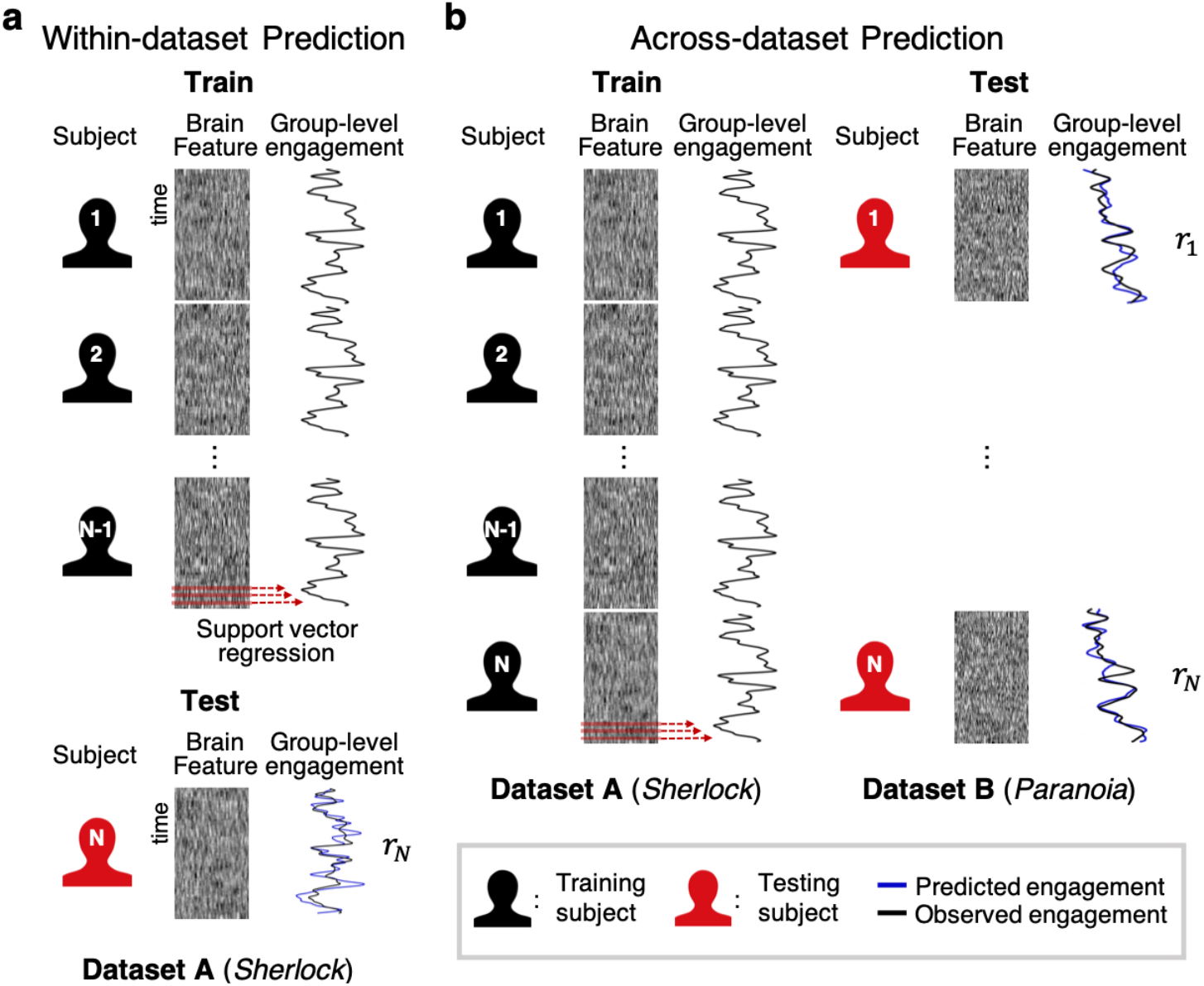
Schematic illustration of dynamic predictive modeling. (**a**) Within-dataset prediction. Internal model validation is conducted using leave-one-subject-out cross-validation. Multivariate patterns of brain activity at time *t* are aligned with a group-level behavioral score (e.g., group-average engagement) at time *t*, across all participants but one. The support vector regression (SVR) model is trained using data from all time points and training participants, and is then applied to a held-out participant’s brain activity time-course to predict engagement at every time point. Prediction accuracy is measured as the average of the cross-validation folds’ Pearson’s correlation (*r*_*N*_) between predicted and observed engagement time-courses. (**b**) Across-dataset prediction. To externally validate predictive models, an SVR is trained using all participants’ brain activity time-courses and the group-level engagement ratings from dataset A (e.g., *Sherlock*). The model is tested on fMRI data from every participant in dataset B (e.g., *Paranoia*). Prediction accuracy is computed by averaging every test participant’s Pearson’s correlation (*r*_*N*_) between predicted and observed behavior time-courses.

We examined whether patterns of BOLD activity in regions selected in the above ISC analysis (i.e., the regions in **Fig. 2b**) predict changing levels of engagement. Models trained on patterns of BOLD signal in these regions did not show robust prediction of group-average engagement ratings (*Paranoia*: *r* = .040, *p* = .468, RMSE = 1.050, *p* = .007, *R*^2^= −.050, *p* = .008; *Sherlock*: *r* = .127, *p* = .037, RMSE = 1.545, *p* = .986, *R*^2^=−.545, *p* = .987). Even when using all 122 Yeo atlas ROIs as features, models did not show robust prediction of engagement (*Paranoia*: *r* = .114, *p* = .263, RMSE = 1.012, *p* = .094, *R*^2^=-.012, *p* = .095; *Sherlock*: *r* = .240, *p* = .004, RMSE = 1.207, *p* = .970, *R*^2^= −.207, *p* = .971). To increase the specificity of the neural features, we conducted additional feature selection in every cross-validation so that only the ROIs of which BOLD time-courses are consistently correlated with group-average engagement were included as features to the model (one-sample t-test, *p* < .01).^47^ Still, BOLD activation patterns did not consistently predict engagement (*Paranoia*: *r* = −.021, *p* = .894, RMSE = 1.120, *p* = .232, *R*^2^= −.120, *p* = .217; *Sherlock*: *r* = .234, *p* < .001, RMSE = 1.223, *p* = .500, *R*^2^= −.223, *p* = .495).

### Functional connectivity predicts changes in engagement, even across different stories

Although multivariate patterns of BOLD activity failed to predict changes in engagement, evidence suggests that changes in the statistical interactions of activity in pairs of brain regions—functional connectivity dynamics—predict cognitive and attentional state changes during task performance.^25,48^ Thus, we examined whether whole-brain FC patterns predict evolving states of engagement.

Time-resolved FC was extracted by computing Pearson’s correlations (Fisher’s *r*-to-*z* transformed) between the time-courses of every pair of ROIs using a tapered sliding window.^37^ Predictive models were first trained and tested within-dataset using LOO cross-validation. SVRs were trained to predict group-average engagement from all but one participants’ multivariate FC patterns, and then applied to dynamic FC patterns from the held-out individual to predict TR-by-TR engagement (**Fig. 3a**). Feature selection was conducted in every round of cross-validation; FC significantly correlated with engagement in the training set were selected as features (one-sample t-test, *p* < .01).^47^ Models predicted group-average engagement ratings above chance for both the *Paranoia* (*r* = .372, *p* = .005; RMSE = .862, *p* = .005; *R*^2^= .138, *p* = .006) and *Sherlock* (*r* = .568, *p* = .005; RMSE = .679, *p* = .005; *R*^2^= .321, *p* = .006) datasets (**Fig. 4a**). Of note, the null distributions are positively skewed (**Fig. 4a**), potentially because SVRs were trained and tested on the same behavioral outcome. Nonetheless, prediction accuracy was significantly above these skewed chance distributions. Predictive power was significantly higher than chance when we used a different type of null distributions generated by randomly selecting the same number of functional connections (FCs) from among those not selected in the actual feature selection (*Paranoia*, *p* < .001, *p* < .001, *p* < .001; *Sherlock*: *p* < .001, *p* < .001, *p* < .001, respectively for Pearson’s *r*, RMSE, and *R*^2^; iteration = 1,000).

**Figure 4.**
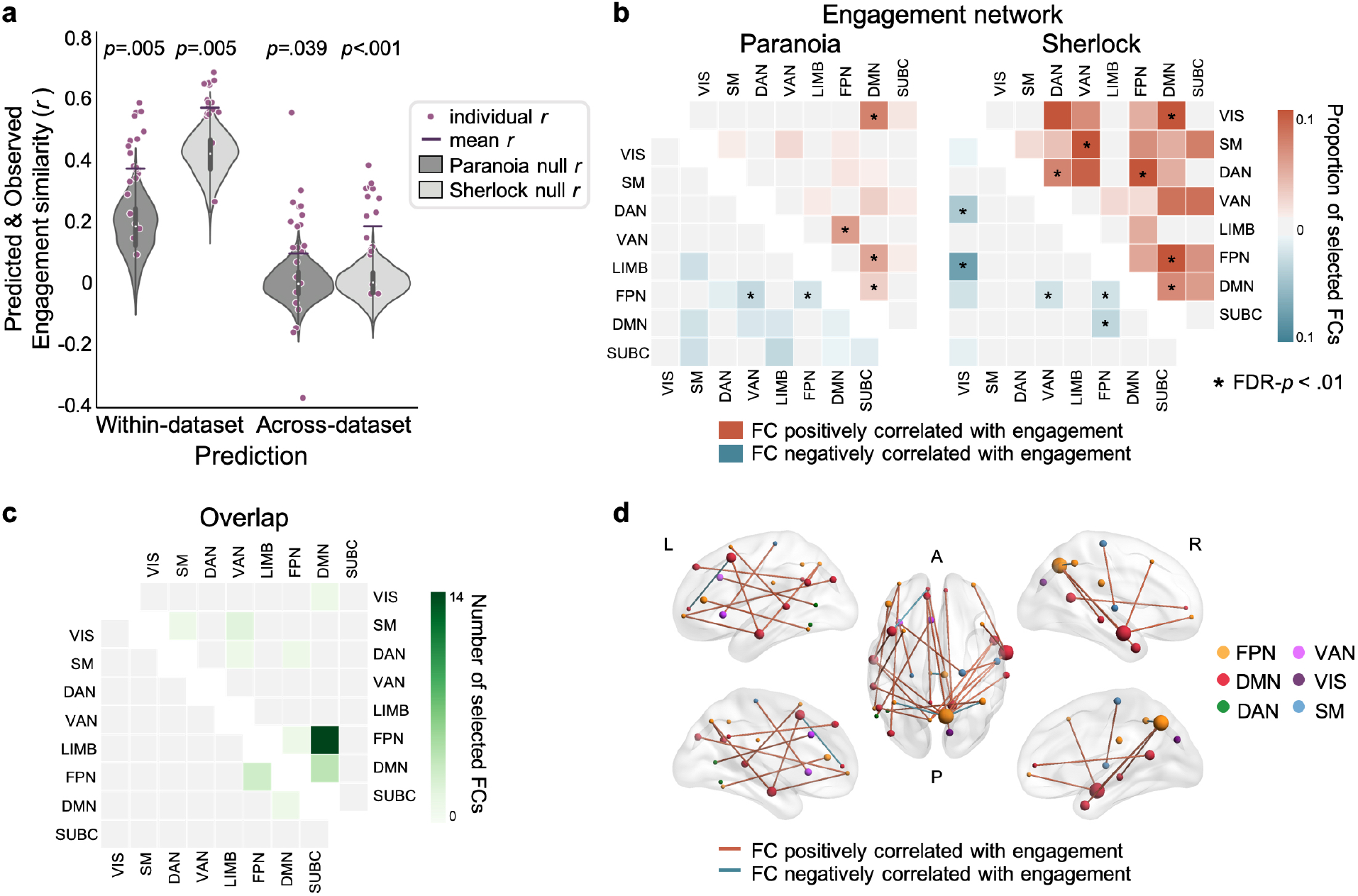
Functional connectivity (FC) predicts group-average, dynamic states of engagement. (**a**) Model performance for the within-dataset (left) and across-dataset (right) predictions for *Paranoia* (dark gray) and *Sherlock* (light gray). Purple dots indicate Pearson’s correlations between predicted and observed engagement time-courses for every participant. Purple bars indicate predictive performance; the mean *r* of the cross-validations. Gray violin plots show null distributions of mean prediction performance, generated from models that predict phase randomized engagement time-courses. The significance of empirical *r* was computed based on the null distribution (one-tailed tests *p* < .05; iteration = 1,000). (**b**) Functional connections (FCs) selected in every cross-validation fold, grouped in pre-defined functional networks.^42^ Colors indicate the proportion of selected FCs from all possible connections of each network pair. The upper triangle matrix (red) illustrates the proportion of FCs positively correlated with engagement, and the lower triangle matrix (blue) illustrates the proportion of FCs negatively correlated with engagement. Network pairs that are selected more than expected by chance are indicated with asterisks (FDR-*p* < .01; iteration = 10,000). (**c**) The number of FCs correlated with engagement dynamics of both the *Paranoia* and *Sherlock*, summarized in functional networks. The upper triangle indicates FCs positively correlated with engagement and the lower triangle indicates FCs negatively correlated with engagement in both samples. (**d**) The overlapping FCs in **c**, visualized in the brain.^49^ Red lines indicate FCs positively correlated with engagement, whereas blue lines indicate FCs negatively correlated with engagement. Node colors indicate different functional networks. Node sizes indicate total number of FCs involving that node. VIS: Visual, SM: Somatosensory motor, DAN: Dorsal attention, VAN: Ventral attention, LIMB: Limbic, FPN: Frontoparietal, DMN: Default mode, SUBC: Subcortical networks.

Within-dataset prediction demonstrates that models based on FC dynamics predict stimulus-related engagement during two distinct narratives. To confirm that models are capturing engagement and not other stimulus-specific regularities, we conducted across-dataset prediction (**Fig. 3b**). SVRs learned the mappings between multivariate patterns of FCs and moment-to-moment engagement using data from all participants in one dataset (e.g., *Sherlock*). The FCs selected in every round of cross-validation during the within-dataset prediction were used as features (**Fig. 4b**). Next, the model trained in one dataset was applied to predict engagement in the held-out dataset (e.g., *Paranoia*). Across-dataset prediction was successful both when predicting engagement of *Paranoia* from a model trained with *Sherlock* (*r* = .096, *p* = .039; RMSE = 1.045, *p* = .042; *R*^2^= −.092, *p* = .042; see **Supplementary Fig. 4** for interpretation of negative *R*^2^) and when predicting engagement of *Sherlock* from a model trained with *Paranoia* (*r* = .185, *p* < .001; RMSE = 1.012, *p* = .002; *R*^2^= −.024, *p* = .002). As expected, the null distribution was not shifted in the across-dataset prediction, which validates that the model did not learn a spurious relationship between FC and story-specific properties other than subjective engagement (**Fig. 4a**). However, for the across-dataset prediction, the predictive power of the engagement networks was not significantly better than that of size-matched random networks selected from FCs outside the engagement networks (*Paranoia*: *p* = .450, *p* = .511, *p* = .512; *Sherlock*: *p* = .002, *p* = .06, *p* = .070, respectively for Pearson’s *r*, RMSE, and *R*^2^). The result implies that narrative engagement may be reflected in widely distributed patterns of time-varying FC, in addition to being specifically predicted by the observed engagement networks.

To characterize the anatomy of the predictive networks, we visualized the FCs consistently selected in every round of within-dataset cross-validation for the *Paranoia* and *Sherlock* datasets (**Fig. 4b**). We call these sets of FCs the *engagement networks*. 205 FCs were consistently selected in *Paranoia*, with 125 FCs positively and 80 FCs negatively correlated with engagement. For *Sherlock*, a set of 675 FCs was selected, with 598 FCs positively and 77 FCs negatively correlated with engagement. A total of 29 FCs was selected in both datasets, with significant degree of overlap (probability of randomness using the hypergeometric cumulative distribution function, *p* = .006). 25 FCs were positively correlated (significance of overlap, *p* < .001) and 4 FCs were negatively correlated (*p* < .001) with engagement in both datasets (visualized in **Figs. 4c** and **4d**).

To examine whether particular functional networks were included in the engagement networks above chance, we calculated the proportion of selected FCs relative to the total number of possible connections between each pair of functional networks (FDR *p* < .01; **Fig. 4b**). Consistently for both datasets, connections between the DMN and visual network, DMN and FPN, as well as within-DMN connections were positively correlated with group-average engagement. Connections between FPN and VAN and within-FPN connections exhibited negative correlations with engagement. The results suggest that the FCs within and between regions of the DMN and FPN contain representations of higher-order cognitive states of engagement that are not specific to sensory modality (visual vs. auditory) or narrative contents (different stories).

### Visual sustained attention network predicts engagement during visual narratives

Previous work theorized that narrative engagement involves attentional focus on story events.^9,10,50,51^ To test this empirically, we asked whether a *sustained attention network*, a set of functional connections that predict individuals’ ability to sustain attention during a gradual-onset CPT (gradCPT),^52^ predicts dynamic changes in engagement during story comprehension. We hypothesized that FCs that predict the ability to stay focused during a controlled experimental task may also reflect the degree to which individuals are engaged in narratives at each moment of time. To do so, we first re-defined the sustained attention network using data from Rosenberg et al.^52^ in Yeo atlas space in order to apply it to the *Paranoia* and *Sherlock* datasets (**Fig. 5a**; see *Methods* for detail). Results revealed that connections between visual, somatosensory-motor, and dorsal and ventral attention networks were positively correlated with individuals’ gradCPT performance, potentially reflecting the fact that the task requires participants to exert goal-directed attention to visually-presented stimuli. On the other hand, connections with the DMN were negatively correlated with gradCPT performance, aligning with conceptions of the DMN as task-negative network in certain contexts.^53,54^

**Figure 5.**
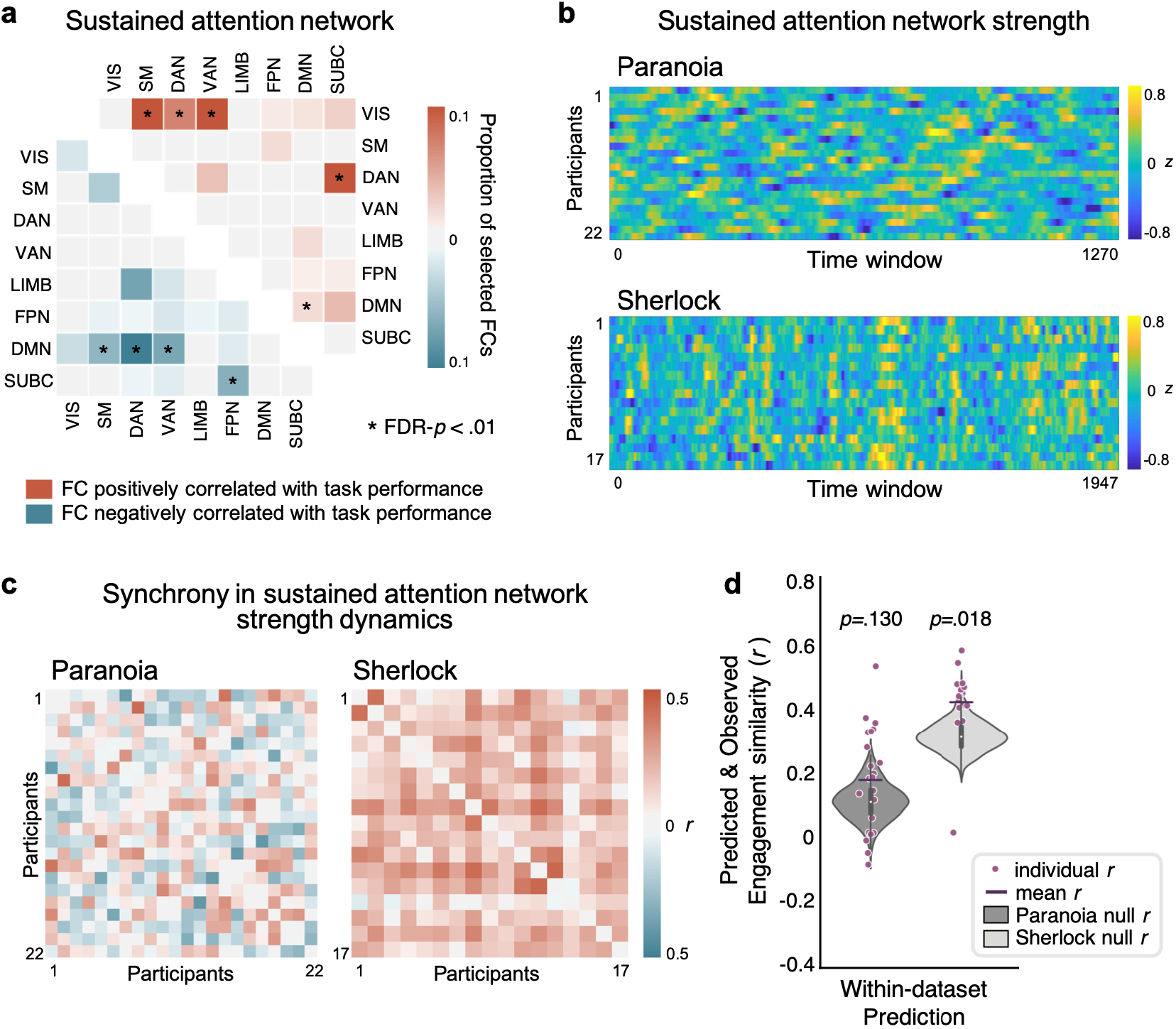
Visual sustained attention network defined from Rosenberg et al.^52^ in relation to group-average engagement. (**a**) Functional connections (FCs) significantly correlated with gradCPT performance of 25 individuals, summarized in Yeo et al.^42^ functional network space. The upper triangle matrix (red) illustrates the proportion of FCs positively correlated with sustained attention scores, and the lower triangle matrix (blue) illustrates the proportion of FCs negatively correlated with sustained attention scores. (**b**) Time-resolved sustained attention network strength of every fMRI participant. Network strength is computed as the difference between the average functional connectivity time-courses of the positively correlated FCs and negatively correlated FCs, z-normalized across time within participant. (**c**) Synchrony (Pearson’s correlation) between pairwise participants’ sustained attention network strength timecourses, illustrated in **b**. (**d**) Predictive performance of the within-dataset prediction, respectively for *Paranoia* (dark gray) and *Sherlock* (light gray). The patterns of FCs included in **a** were used as multivariate features in the model.

To test whether sustained attention network dynamics were time-locked to the narratives, we examined whether network strength was synchronized across individuals comprehending the same story. **Fig. 5b** depicts the time-resolved network strength of all participants. For *Paranoia*, pairwise subject similarity was, on average, *r* = .002, with only 38.10% of pairwise participants showing positive correlations (FDR-*p* < .05; **Fig. 5c**). For *Sherlock*, however, participants exhibited significant degrees of synchrony in sustained attention network strength, with a mean *r* = .182 and 89.71% of pairwise participants showing positive correlations (FDR-*p* < .05; **Fig. 5c**). We examined whether the dynamics of sustained attention network strength are correlated with group-average engagement. We did not observe evidence of correlations between sustained attention network dynamics and engagement (mean of the correlations between each participant’s sustained attention network strength time-course and the group-average engagement time-course: *Paranoia*: *r* = −.004, *p* = .900; *Sherlock*: *r* = .052, *p* = .475), suggesting that the sustained attention network does not explicitly represent fluctuating states of narrative engagement in its strength alone (but see **Supplementary Fig. 5** for correlation between dynamic ISC, a brain-based measure of stimulus-related activity, and averaged sustained attention network strength).

We next asked if multivariate patterns of time-resolved FCs in the sustained attention network predict group-average engagement (**Fig. 5d**). Indeed, when trained with LOO cross-validation, SVRs based on FC patterns of the sustained attention network predicted changes in engagement in the *Sherlock* dataset (*r* = .434, *p* = .018, RMSE = .816, *p* = .020, *R*^2^= .184, *p* = .021), in which the narrative was delivered in the auditory and visual modalities, but not in the *Paranoia* dataset (*r* = .189, *p* = .130, RMSE = .996, *p* = .158, *R*^2^= .004, *p* = .159) which was delivered only in the auditory modality. Predictive power was robust for *Sherlock* when compared to the size-matched random networks selected from FCs outside the engagement network and sustained attention network (*p* = .035, *p* = .042, *p* = .043, respectively for Pearson’s *r*, RMSE and *R*^2^). The results provide empirical evidence that the sustained attention network—which was defined in a completely independent study using a controlled experimental paradigm requiring top-down attentional control—predicts stimulus-related engagement during story comprehension.

Finally, we characterized the anatomical overlap between the engagement network of *Sherlock* (**Fig. 4b** right) and the sustained attention network (**Fig. 5a**). There was a significant overlap of the FCs that positively correlated with both sustained attention and engagement of *Sherlock* (calculated using the hypergeometric cumulative distribution function, *p* = .002). Connections between the visual network and the attention networks (dorsal and ventral), and connections between the DMN and FPN were included in both networks above chance (FDR *p* < .05). On the other hand, we observed no significant overlap of the negatively correlated FCs (*p* = .881). The results were robust when we calculated anatomical overlap using a range of feature-selection thresholds (**Supplementary Table 3**). The partial overlap between the networks predicting sustained attention and engagement suggests that common processes are involved in maintaining focus on a visual sustained attention task and a visual narrative.

### Engagement network during memory encoding predicts later recall of the events

During both a television episode and an audio-narrated story, people’s engagement fluctuated over time with patterns of functional brain connectivity. What consequences do these changes in narrative engagement have for how we remember stories? To address this question, we asked whether attentional engagement facilitates encoding of events into long-term memory. To measure how well each moment of the stories was remembered, we analyzed fMRI participants’ free recall data provided by Finn et al.^18^ and Chen et al.^13^ Again, to test for the robustness of results with different analysis pipelines, we used two different approaches to quantify recall. For *Paranoia*, each phrase of participants’ recall was manually matched to the semantically closest sentence in the original story transcript (**Fig. 6a**). Every word in the story and recall transcripts was then represented as a vector in a distributional word embedding space, GloVe,^55^ and the degree of similarity between the matched recall and transcript was computed using the cosine similarities between the bag-of-words embedding vectors.^56^ For *Sherlock*, we used a data-driven topic modeling approach^57^ to segment the stories into multiple events and quantified the similarities between recalled events with the video annotations (**Fig. 6b**). **Fig. 6c** illustrates exemplars of different metrics that represent subsequent “recall fidelity” time-course.

**Figure 6.**
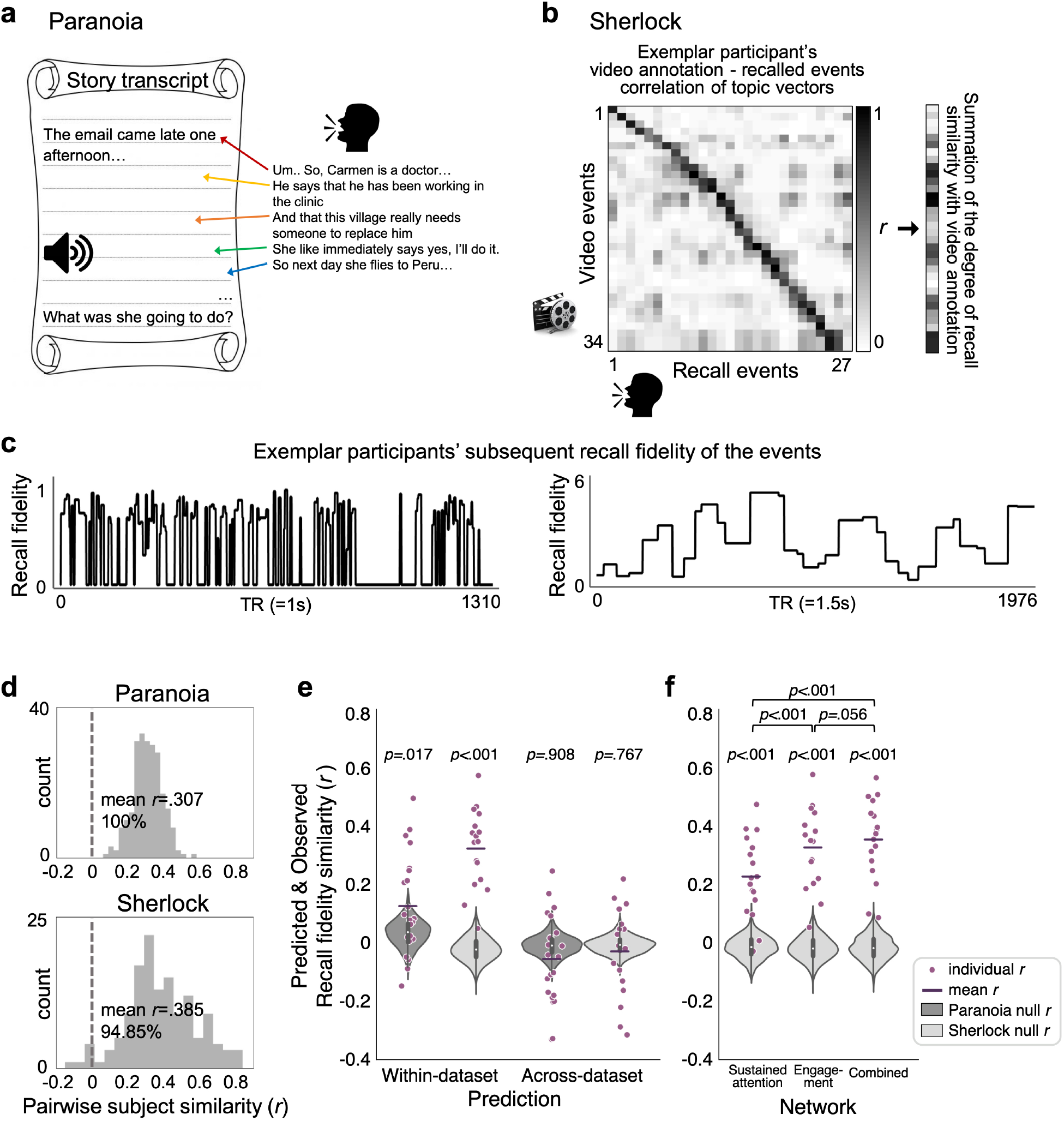
Individual-specific story recall in relation to group-average engagement. To quantify the precision with which events in a story were recalled, (**a**) in *Paranoia*, we matched the phrases uttered during free recall to the story transcript, and calculated semantic similarity using distributional word embeddings. (**b**) In *Sherlock*, we applied dynamic topic modeling fitted with video annotations to extract topic vectors of individuals’ recall transcripts. The topic vector similarities between recall and video annotations were summed across recalled events.^57^ (**c**) Representative participants’ subsequent recall fidelity of the events for *Paranoia* (left) and *Sherlock* (right). (**d**) Histograms of the pairwise subjects’ recall similarity values. Recall similarity was calculated as the Pearson’s correlation between pairs of participants’ recall fidelity time-courses. (**e**) Predictive performance of individual-specific recall fidelity from the engagement networks of within-dataset (left) and across-dataset (right) for *Paranoia* (dark gray) and *Sherlock* (light gray). The patterns of FCs that predicted group-average engagement, illustrated in **Fig. 4b**, were used as multivariate features in the model. (**f**) Predictive performance of individual-specific recall fidelity of *Sherlock* dataset using different network features; sustained attention network (left), engagement network (middle), and the combined network which contains FCs that are either predictive of engagement or sustained attention. The upper brackets indicate significance of the comparisons between the pairwise model performances (paired t-tests between the cross-validation performances).

To examine whether people tend to remember and forget similar events, we calculated the similarity of recall fidelity dynamics of all pairwise fMRI participants (**Fig. 6d**). Almost every pairwise participants exhibited similar trends of recall, for both *Paranoia* (mean *r* = .307, with 100% of pairwise participants showing positive correlations, FDR-*p* < .05) and *Sherlock* (mean *r* = .385, with 94.85% showing positive correlations, FDR-*p* < .05). The results imply that particular events in a story tend to be more likely to be recalled than other events, replicating previous empirical findings of Meyer and McConkie,^58^ and revisiting theoretical explanations that the subjective importance of events within situational contexts influences memory for those events.^59^

We asked whether fMRI participants’ individual-specific recall scores were significantly related with behavioral participants’ group-average engagement. Correlations between individual-specific recall time-courses and engagement were averaged, and then compared with the null distribution where engagement ratings were phase randomized (iteration = 10,000). We observed different trends of results for the two story datasets: significant correlations between engagement and recall in *Sherlock* (average *r* = .328, *p* = .007), but no relationship observed in *Paranoia* (*r* = .016, *p* = .804).

The disparate results may be due to story-specific components, differences in the behavioral analysis pipelines, or a possibility that behavioral engagement ratings do not fully capture attention fluctuations that are consequential for memory. Therefore, we applied a dynamic predictive modeling approach, asking whether each narrative’s engagement network (**Fig. 4b**) predicted corresponding narrative’s individual-specific story recall (**Fig. 6c**). Models predicted the fidelity to which each moment of the story would be later recalled (**Fig. 6e;**replacing group-level engagement to individual-specific recalls in **Fig. 3**) in both the *Paranoia* (*r* = .135, *p* = .017; RMSE = 1.043, *p* = .015; *R*^2^= −.032, *p* = .016) and *Sherlock* datasets (*r* = .334, *p* < .001; RMSE = .901, *p* < 0.001; *R*^2^= .099, *p* < .001). Furthermore, demonstrating specificity, the engagement network better-predicted recall than did size-matched random networks selected from FCs outside the engagement networks; *Paranoia*: *p* = .049, *p* = .099, *p* = .101; *Sherlock*: *p* = .002, *p* < .001, *p* = .002, respectively for Pearson’s *r*, RMSE and *R*^2^). However, models did not generalize to predict recall across datasets (*Paranoia* predicted from a model trained with *Sherlock*: *r* = −.048, *p* = .908; RMSE = 1.206, *p* = .999; *R*^2^= −.206, *p* = 1.0; *Sherlock* predicted from a model trained with *Paranoia*: *r* = −.021, *p* = .767; RMSE = 1.115, *p* = .479; *R*^2^= −.115, *p* = .480).

Finally, we examined whether FCs in the sustained attention network (**Fig. 5a**) also predict individual-specific story recall of the *Sherlock* sample (**Fig. 6f**). (We did not test whether sustained attention network FCs predicted recall in the *Paranoia* dataset because they did not predict engagement in that sample.) Sustained attention network connections predicted subsequent memory of *Sherlock* (*r* = .233, *p* < .001, RMSE = .979, *p* < .001, *R*^2^ = .021, *p* < .001), but with less predictive power than the engagement network model (paired t-test, *t*(16) = 4.451, *p* < .001). This result suggests that even though the two networks were defined in different contexts (i.e., top-down attentional control or stimulus-related narrative engagement), both may share a role in predicting subsequent memory. A combined network model including FCs that predict engagement and sustained attention also predicted recall above chance (*r* = .362, *p* < .001, RMSE = .876, *p* < .001, *R*^2^ = .124, *p* < .001). However, the combined network model did not explain additional variance in subsequent memory compared with the engagement network model alone (*t*(16) = 2.063, *p* = .056). Thus, sustained attention network FCs explain only a subset of the variance in subsequent event memory captured by engagement network FCs.

## Discussion

The degree to which we are engaged in narratives in the real world fluctuates over time. Sometimes we may find ourselves immersed in a particularly engrossing story, whereas at other moments we may become bored with a plot line or struggle to follow a conversation thread. What drives these changes in narrative engagement, and how does engagement affect what we later remember about a story?

Here, using data from independent behavioral and fMRI studies, we test theoretical proposals that narrative engagement reflects states of heightened attentional focus and emotional arousal. Specifically, we analyzed two open-source fMRI datasets collected as participants watched an episode of *Sherlock* or listened to an audio-narrated story, *Paranoia*. We ran behavioral experiments in which independent participants continuously rated how engaging they found the narratives. Self-reported engagement fluctuated across time and was synchronous across individuals, demonstrating that this paradigm captures states of engagement that are shared across individuals (**Fig. 1**). Group-average changes in engagement were correlated with the narratives’ emotional contents, but not their low-level sensory salience. Furthermore, dynamic ISC revealed that stimulus-evoked BOLD responses in regions of the DMN scaled with group-average engagement, suggesting that DMN activity becomes *more synchronized* across individuals when people are *more engaged* in the narratives (**Fig. 2**). Fully cross-validated models (**Fig. 3**) trained on time-resolved whole-brain FC, but not BOLD activity, predicted moment-to-moment changes in engagement in each fMRI sample. Predictive models further generalized across datasets (**Fig. 4**), demonstrating that FC contains information about higher-order, modality- and context-general cognitive states of engagement. We observed that the sustained attention network, defined using data collected during a visual CPT, predicted changes in engagement as individuals watched *Sherlock*, but not as individuals listened to *Paranoia* (**Fig. 5**), suggesting that brain networks involved in top-down attentional control may be partially involved when individuals are attentively engaged in naturalistic contexts. Lastly, we found that the engagement network, as well as the sustained attention network, predicted how likely participants were to remember certain story events (**Fig. 6**), suggesting that dynamic attentional engagement during encoding affects later memory for narrative events.

Historically, the majority of research on sustained attention has studied attention to external stimuli during psychological tasks requiring top-down attentional control and cognitive effort.^5,60–62^ Complementary work has described states of attention associated with less subjective effort, such as *flow* states of complete absorption in an activity,^8^ *in-the-zone* moments of optimal task performance,^63–66^ and *soft fascination* to natural environments or aesthetic pieces of art.^67,68^ Recent work has also characterized stimulus-independent internal attention and mind wandering.^7,69–72^ However, relatively few studies have tackled attentional mechanisms during the processing of contextually-rich, everyday narratives to ask, for example, to what degree narrative engagement requires attentional control, how attentional fluctuations modulate moment-to-moment narrative comprehension and memory, or how state-like or even trait-like differences in subjective engagement are reflected in the brain activity. Bellana and Honey^73^ theorized that *deep information processing*^74^ is involved when comprehending narratives, which leads to high engagement (or transportation, absorption, and immersion) during perception and rich memory representation. Our work provides empirical support that the neural signatures involved in top-down sustained attention (**Fig. 5a**) and stimulus-related narrative engagement (**Fig. 4b**) are partially overlapping, and that these two networks both play a role in predicting subjective engagement and subsequent memory. This study aims to bridge the gap in the literature, asking to what degree our current knowledge of attention and memory in controlled experimental settings explains real-world cognition. Future studies should aim to characterize the common and distinct components of attentional processes and neural signatures in multiple situational contexts.

Previous research has studied interactions between sustained attention and subsequent memory using controlled psychological tasks that measure participants’ memory for stimuli encountered in different attentional states. For example, participants in a study from deBettencourt et al.^4^ performed a CPT and then reported their incidental recognition memory of the presented images. CPT stimuli seen during engaged attentional states (indexed with response times) were better remembered. Complementary work examined relationships between sustained attention and recall during naturalistic task using an individual differences approach. Jangraw et al.^75^ asked participants to read transcripts of Greek history lectures during fMRI, then measured their performance on a post-scan comprehension test. Individuals who showed FC signatures of stronger sustained attention during reading performed better on comprehension test. However, attending to and remembering events constantly fluctuates across time, heavily dependent on the situational contexts and relational structure. It is yet unknown the role of dynamic memory when constructing structured representation of narratives,^76^ or how we modulate attention online to encode critical events and reinstate relevant memories. The implication of our results—that neural signatures of engagement predict subsequent memory—motivate future work addressing neural mechanisms of the complex, bidirectional interactions between sustained attention and memory in naturalistic contexts.

The current study focuses on group-level states of engagement that are shared across individuals and are thus stimulus-related. Future work characterizing each person’s unique pattern of attentional engagement during narratives—perhaps assessed with physiological measures such as heart rate, electrodermal activity,^77,78^ or facial expressions,^79^ or with pupil dilation^80^ and blink rate^81,82^—can help disentangle intrinsic from stimulus-related attention fluctuations and elucidate their consequences for memory at the individual level.

In sum, we show that engagement during story comprehension dynamically fluctuates across time, driven by narrative content. The study characterizes relationship between sustained attention and narrative engagement, and elucidates neural signatures that predict future event memory.

## Acknowledgements

We thank Janice Chen and colleagues for open-sourcing the *Sherlock* dataset and Andrew C. Heusser and colleagues for sharing code on topic modeling. We thank Jeongjun Park for help with conceptualization. Our work was supported by the University of Chicago Social Sciences Division, resources provided by the University of Chicago Research Computing Center, and a Neubauer Family Foundation Distinguished Scholar Doctoral Fellowship from the University of Chicago to H.S.

## Author Contributions

Conceptualization, H.S., E.S.F. and M.D.R; Methodology, H.S., and M.D.R.; Formal Analysis, H.S.; Investigation, H.S., E.S.F. and M.D.R; Writing - Original Draft, H.S.; Writing - Review & Editing, H.S., E.S.F. and M.D.R.; Supervision, M.D.R.

## Methods

### Behavioral Experiments

21 individuals from the University of Chicago or the surrounding community participated in a behavioral study in which they listened to the *Paranoia* story stimulus (3 left-handed, 11 females, mean age 23.57 ± 4.04; 12 native English-speakers, 2 bilinguals). 17 individuals participated in a behavioral study in which they watched an episode of television series *Sherlock* (1 left-handed, 11 females, mean age 22.06 ± 3.75; 11 native English-speakers, 2 bilinguals). All participants reported no history of visual or hearing impairments, provided written informed consent before the study, and were compensated for their participation. The study was approved by the Institutional Review Board of the University of Chicago.

Stimulus presentation and response recording were controlled with PsychoPy3.^83^ The experiment took place in a dimly lit room in front of a Macintosh Apple monitor. The participants either listened to a 20 min audio-narrated *Paranoia* or watched a 50 min episode of *Sherlock*, while continuously rating how engaging they found the story by adjusting a scale bar from 1 (“Not engaging at all”) to 9 (“Completely engaging”). The scale bar was constantly up on the bottom of the computer monitor. Other aspects of the experiment tried to replicate, as closely as possible, previous fMRI studies: the same visual image was shown at the center of the screen as participants listened to *Paranoia*,^18^ and “run breaks” occurred at exactly the same moments for both stories.

To confirm participant compliance, participants who listened to *Paranoia* completed nine comprehension questions asking about the events. The average percent correct was 95.24 ± 8.29%, verifying that all participants maintained overall attention to the story. The participants who watched *Sherlock* were asked to provide a spoken recall of the story in as much detail as they remember as soon as they finished watching the episode. All participants were qualitatively assessed to have paid attention to the story.

### Functional MRI Image Acquisition and Preprocessing

The raw structural and functional images of the *Paranoia* dataset were downloaded from OpenNeuro (https://openneuro.org/datasets/ds001338/). Functional images of Finn et al.^18^ were acquired from 3T Siemens TrimTrio, using T2+-weighted echo planar imaging (EPI) multiband sequence (TR = 1000 ms, TE = 30 ms, voxel size = 2.0 mm isotropic, flip angle = 60°, field of view = 220 mm × 220 mm). Structural images were bias-field corrected and spatially normalized to the Montreal Neurological Institute (MNI) space. Functional images were motion-corrected using six rigid-body transformation parameters and registered to MNI-aligned T1-weighted images. White matter and CSF masks were defined in MNI space and warped into the native space. Linear drift, 24-parameter motion parameters, mean signals from CSF and white matter, and mean global signal were regressed from the BOLD time-course. We applied a band-pass filter (0.009 Hz < *f* < 0.08 Hz) to remove low frequency confounds and high frequency physiological noise. The data were spatially smoothed with a Gaussian kernel of full width at half maximum (FWHM) of 4 mm.

The preprocessed images of *Sherlock* dataset were downloaded from URL (http://dataspace.princeton.edu/jspui/handle/88435/dsp01nz8062179). Functional images of Chen et al.^13^ were collected from 3T Siemens Skyra, using T2+-weighted EPI sequence (TR = 1500 ms, TE = 28 ms, voxel size = 3.0 × 3.0 × 4.0 mm, flip angle = 64°, field of view = 192 mm × 192 mm). Preprocessing steps followed slice timing correction, motion correction, linear detrending, high-pass filtering (140 s cutoff), coregistration, and affine transformation to the MNI space. The functional images were resampled to 3 mm isotropic voxels.

All analyses were conducted in the volumetric space using AFNI. The cortical surface of the MNI standard template was reconstructed using Freesurfer^84^ for visualization purposes.

### Whole-Brain Parcellation

Cortical regions were parcellated into 114 ROIs following Yeo et al.^41^ based on a seven-network cortical parcellation estimated from the resting-state functional data of 1,000 adults.^42^ Subcortical regions were parcellated into eight ROIs, corresponding to the bilateral amygdala, hippocampus, thalamus, and striatum, extracted from the Freesurfer segmentation of the FSL MNI152 template brain.^41^ The time-courses of the voxels corresponding to each of the 122 ROI were averaged to a single, representative time-course.^25^ The eight functional networks include visual (VIS) and somatosensory-motor networks (SM) relevant to sensory-motor processing, dorsal and ventral attention networks (DAN and VAN) relevant to top-down guided attention or attentional shifts,^85^ default mode (DMN) and frontoparietal networks (FPN) relevant to transmodal information processing,^86^ and the subcortical network (SUBC). To examine the robustness of our results using different functional brain atlas, we used Shen atlas^43^ which parcellates the whole-brain into 268 ROIs, including cerebellum and brainstem in addition to the cortical and subcortical regions.

### Dynamic inter-subject correlation (ISC)

A tapered sliding window was applied to calculate pairwise subjects’ BOLD time-course ISC. We applied the same hyperparameters as when calculating time-resolved FC: a sliding window size of TR = 40 (= 40s) for *Paranoia* and TR = 30 (= 45s) for *Sherlock* datasets, with a step size of 1TR and a Gaussian kernel σ = 3TR. Non-parametric permutation tests were used to quantify the significance of the dynamic ISC of each ROI. Null distributions were generated by taking Pearson’s *r* between the actual dynamic ISC and the phase randomized engagement ratings (two-tailed *t*-test, uncorrected for multiple comparisons, iteration = 10,000).

We tested whether certain functional networks are more likely to be selected as significant in each dataset, as well as for the overlapping regions. We randomly selected the same number of regions in each dataset (21 among 122, and 20 among 122 regions), and calculated the number of regions that correspond to each of the eight pre-defined functional networks. The number and functional network indices of the overlapping regions were also calculated from the surrogate selections. The actual values were compared with the null distributions using a one-tailed test (iteration = 100,000).

### Dynamic predictive modeling

For models based on FC patterns, time-resolved FC matrices were computed as the Pearson’s correlations (Fisher’s *r*-to-*z* transformed) between the BOLD signal time-courses of every pair of regions (122 × 122 ROIs) using a tapered sliding window. SVR models were implemented with python (sklearn.svm.SVR; maximum iteration set to 1,000 and using rbf kernel). The time series of every feature was *z*-normalized across time to retain temporal variance while ruling out individual variance. Every time step was treated as an independent sample.

In addition to measuring prediction performance represented with a mean *r* across cross-validation folds, we tested prediction performance respectively for each cross-validation fold (**Supplementary Figs. 6 and 7**). Almost every cross-validation exhibited higher correlation between predicted and observed engagement over chance, indicating the robustness of the model despite the small number of samples.

**Fig. 4b** and **Fig. 5a** illustrate pairwise regions that were selected in every cross-validation fold, grouped by the ROI’s pre-defined functional network. The proportion of selected pairs was compared with chance using non-parametric permutation test. The empirical proportion was compared with a null distribution in which the same number of features was randomly selected from all possible pairs of regions (FDR-*p* < .01, corrected for network pairs; iteration = 10,000).

### Sustained attention network

We re-analyzed raw fMRI data from Rosenberg et al.,^52^ which were collected from 25 participants as they completed 3 runs—4 blocks, with each 3 min duration—of a sustained attention task, the gradCPT.^63^ Functional image preprocessing steps matched those applied to *Paranoia* data (and differed from those applied to *Sherlock* data). We used Yeo et al.’s^42^ 122 ROI parcellation scheme to directly compare with our results and increase the computational feasibility of dynamic FC pattern-based predictive modeling (the Yeo atlas has 122 ROIs, whereas the Shen atlas, which was originally used in Rosenberg et al.^52^, has 268). The behavioral performance of each gradCPT block was assessed with sensitivity (*d’*). For each participant, overall task performance was computed by taking the average *d’* of all task blocks.

Replicating Rosenberg et al.^52^ with Yeo atlas ROIs, we employed Connectome Predictive Modeling (CPM)^47^ to predict *d’* scores. In a LOO cross-validation approach, we selected FCs that were significantly correlated with training participants’ *d’* scores (Spearman’s correlation, *p* < .01). We separated the FCs that were positively or negatively correlated with *d’*, and averaged the strength of positively correlated FCs and negatively correlated FCs. A multiple linear regression model was trained to learn the coefficients, where the dependent variable was individuals’ *d’* scores, and the two independent variables were the average of positively and negatively correlated FCs, respectively. The model predicted held-out participant’s *d’* score, and the predicted scores of all cross-validations were validated by taking the Pearson’s *r* with the observed scores. Replicating findings from Rosenberg et al.,^52^ FC matrices using Yeo atlas significantly predicted individual’s sustained attention scores (*r* = .801, *p* < .001; comparable to *r* = .822, *p* < .001 when using Shen atlas’ 268-ROI parcellation). The prediction was robust even when we predicted held-out participants’ *d’* scores of individual runs (*r* = .780, *p* < .001), or individual blocks of every run (*r* = .495, *p* < .001).

The FCs selected in every cross-validation are collectively termed the *sustained attention network*. Dynamic strength of sustained attention network was computed by taking the difference between average FC time-courses of positively correlated FCs and the negatively correlated FCs, calculated with the same sliding window analysis as above.

### Event recall

For *Paranoia*, each phrase of a participant’s recall was manually matched to a sentence in the story transcript, which was annotated with timestamps of when that sentence was heard by participants during fMRI. We vectorized all words included in the transcript and recall data with the pre-defined, distributed word embedding, GloVe, which is trained on Wikipedia and Gigaword 5 corpus (dimension = 100).^55^ The event embeddings were generated by the bag-of-words (i.e., average) of the embedding vectors of every word included in the recalled phrase or a sentence in a transcript. The cosine similarity between the event embeddings was multiplied by the binary index of whether a participant recalled the event or not. The output recall scores were extended in time so as to match the moments when a sentence was uttered in the narration.

For *Sherlock*, we applied dynamic topic modeling^87^ combined with hidden Markov model,^11^ implemented by Heusser et al.^57^ We followed the hyperparameter selections and experimental steps of Heusser et al.^57^ We calculated the degree of similarities of recall events with all possible annotated events. The recall fidelity was computed by summing the similarities of recall with every video event.

## Supplementary Information

**Supplementary Figure 1.**
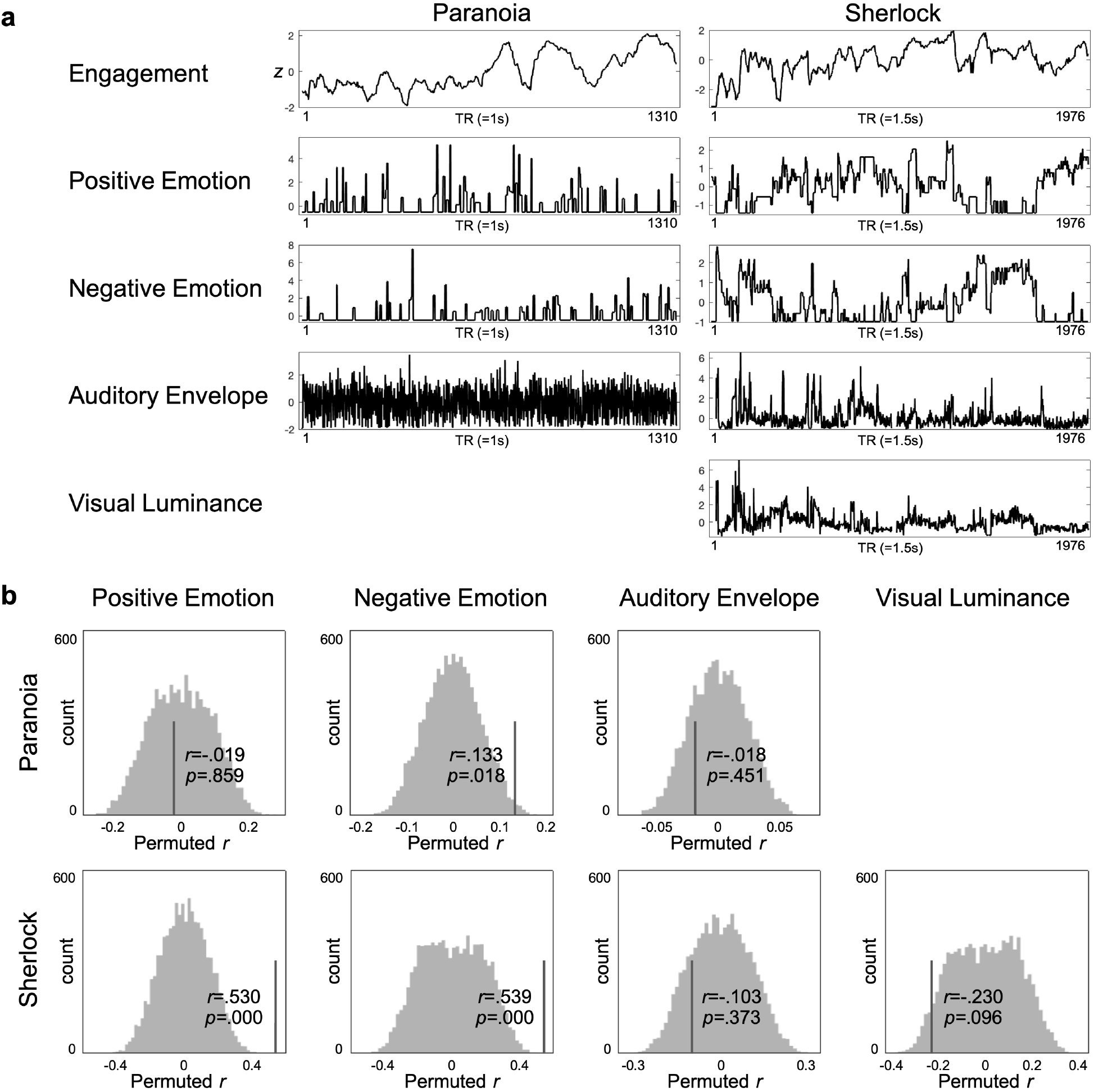
Relationships between attentional engagement and multiple features of the narratives. (**a**) Time-courses of group-average engagement and four story components of *Paranoia* (left) and *Sherlock* (bottom). The time-courses were *z*-normalized across time. (**b**) Results of partial correlations between group-average engagement and each of the four story components, controlling for the other components, from the *Paranoia* (top) and *Sherlock* (bottom) datasets. Histograms indicate distributions of the null partial correlations with phase randomized engagement ratings (iterations = 10,000). The lines indicate empirical partial correlation values, with the *p*-values indicating non-parametric, two-tailed significance tests.

**Supplementary Figure 2.**
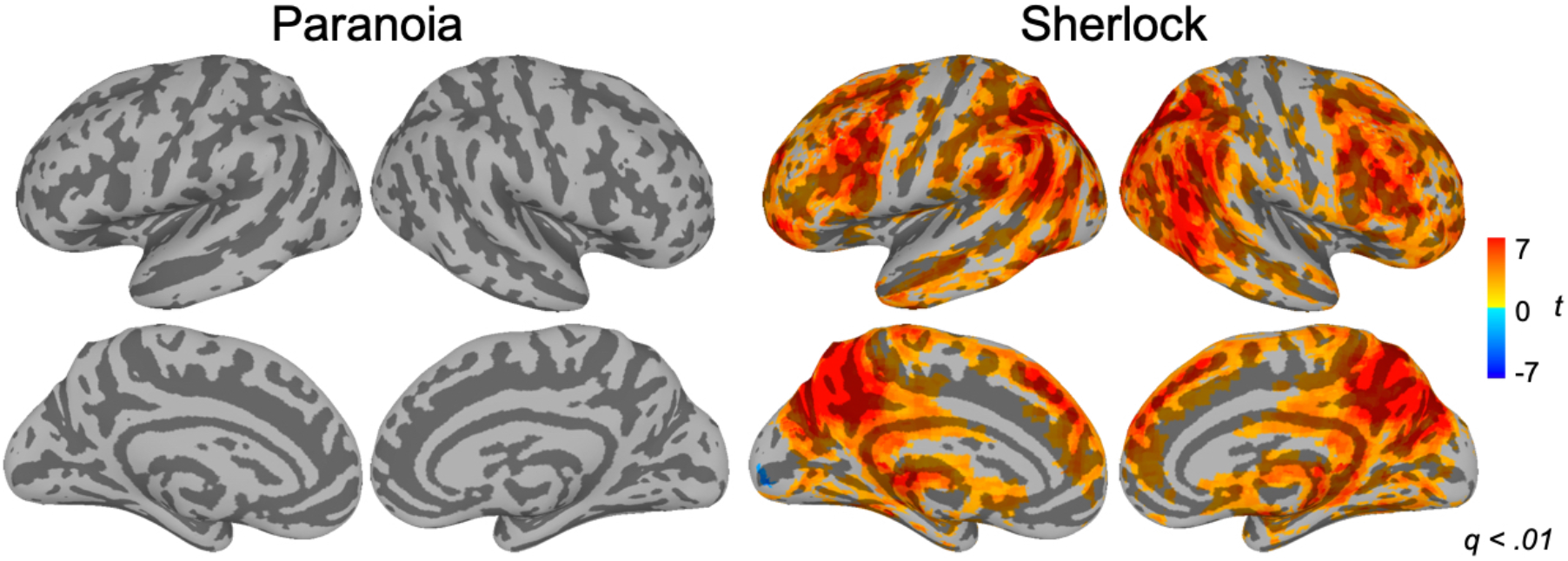
General linear regression model (GLM) results relating group-average engagement time-courses to BOLD activity during *Paranoia* (left) and *Sherlock* (right) (*z* = 2.58 from *q* < .01; cluster size = 35 and 44 voxels respectively, estimated from 3dFWHMx and 3dClustSim using AFNI; individual-voxel *p* < .001, cluster-corrected α < .05). The auditory envelope and visual luminance were controlled for in the GLM. No region’s activity was modulated by engagement in *Paranoia*, whereas activity in almost every region, except somatosensory-motor and early visual and auditory regions, positively scaled with changes in *Sherlock* engagement.

**Supplementary Figure 3.**
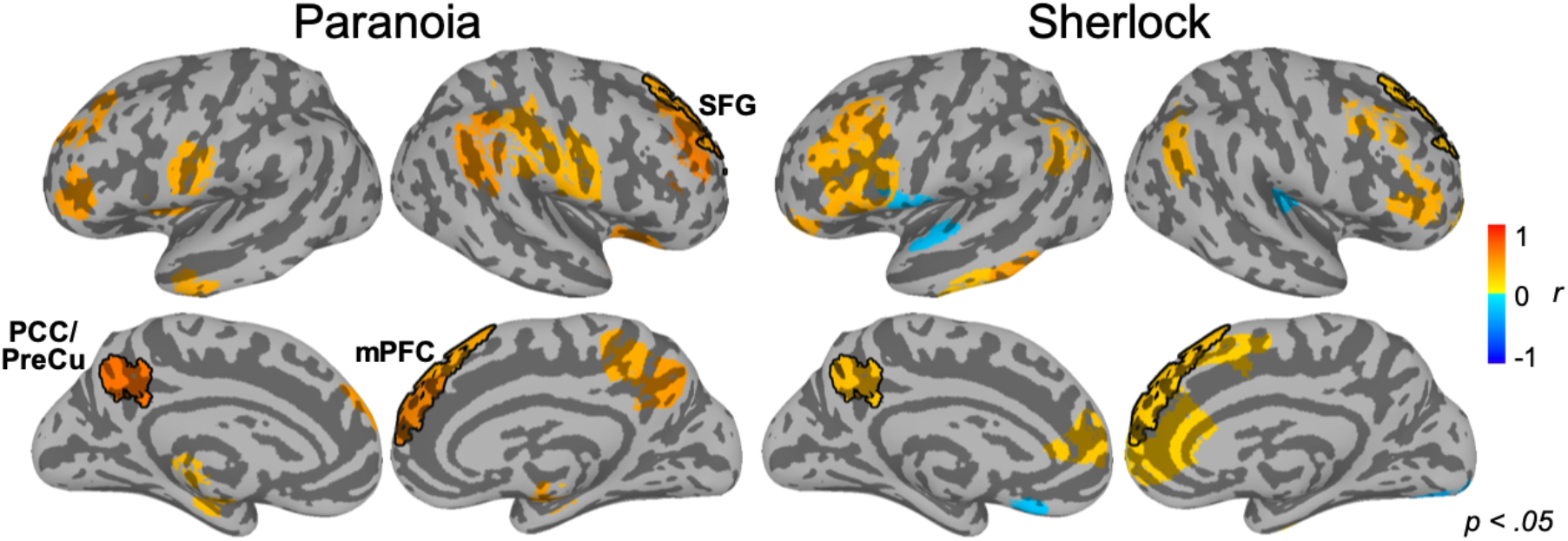
Regions that show significant correlations between dynamic ISC and group-average engagement, using 268-ROIs Shen atlas (two-tailed test; uncorrected *p* < .05). For both datasets, 29 out of 268 regions survived significance tests, with 29/29 regions of *Paranoia* and 25/29 regions of *Sherlock* datasets exhibiting positive correlations between ISC and engagement. The ISCs of the four regions–right superior medial gyrus [−8.5, −53.3, +23.9] (*r* = .562, *r* = .281 for *Paranoia* and *Sherlock* respectively), right superior frontal gyrus [−14.6, − 36.7, +49.1] (*r* = .421, *r* = .304), right caudate [12.8, −13.0, +11.3] (*r* = .459, *r* = .250), and left precuneus [+6.4, +54.3, +37.4] (*r* = .653, *r* = .341)–exhibited significant positive correlations with group-average engagement in both datasets. PCC/PreCu: posterior cingulate cortex/precuneus, mPFC: medial prefrontal cortex, SFG: superior frontal gyrus.

**Supplementary Figure 4.**
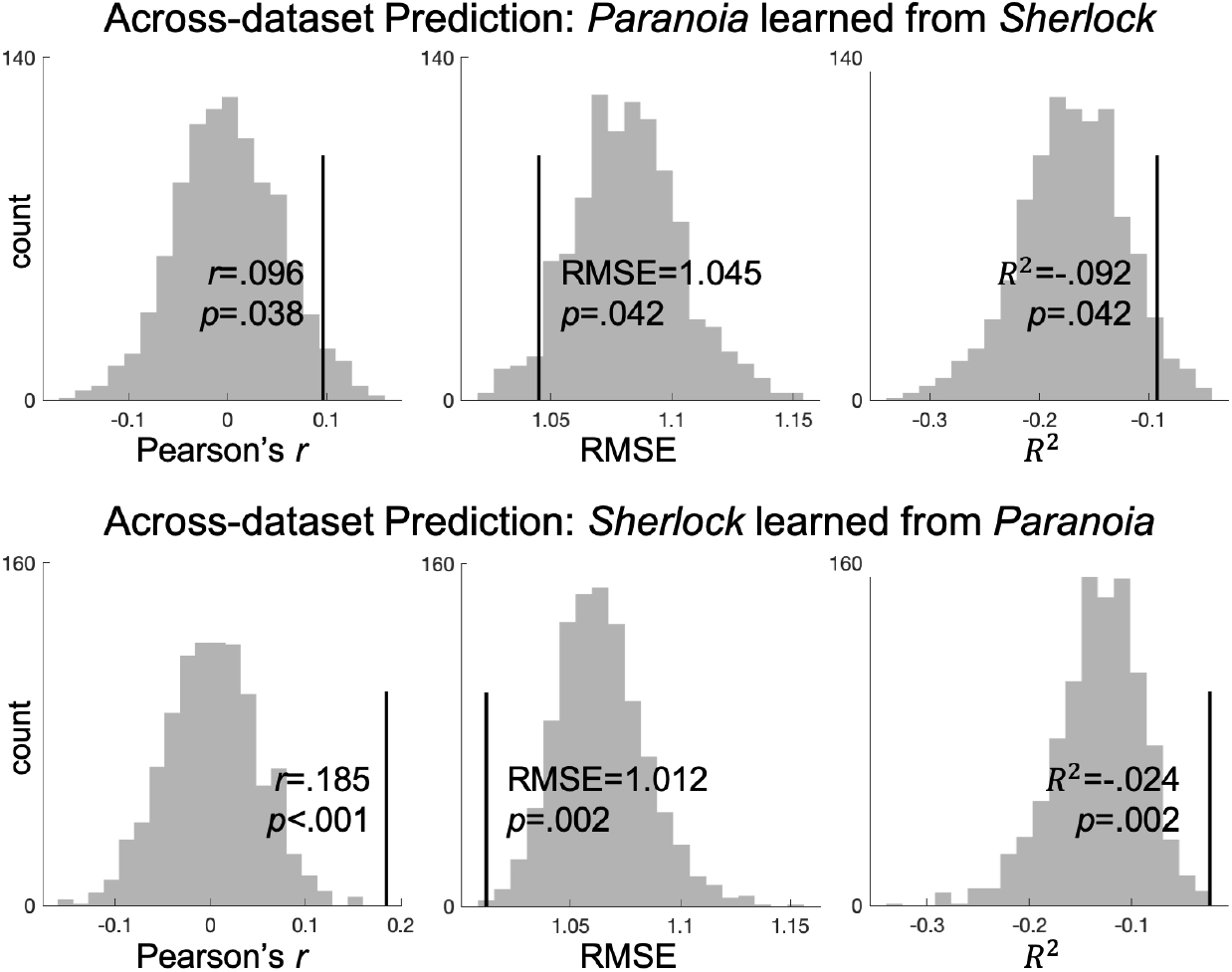
Across-dataset prediction of group-average engagement from dynamic patterns of functional connectivity. The histograms indicate Pearson’s *r* (left), RMSE (center), and *R*^2^ (right) values of null model performance that predicted phase randomized engagement ratings (iteration = 1,000). The empirical values are indicated with vertical lines, with *p* values indicating one-tailed significance tests. Notably, both the null and empirical *R*^2^ values are negative, and the empirical *R*^2^ is consistently higher than chance. These results support our choice of Pearson’s correlation between predicted and observed time-courses as an indicator of prediction performance, reflecting the degree to which models capture temporal dynamics rather than actual values at each moment of time.

**Supplementary Figure 5.**
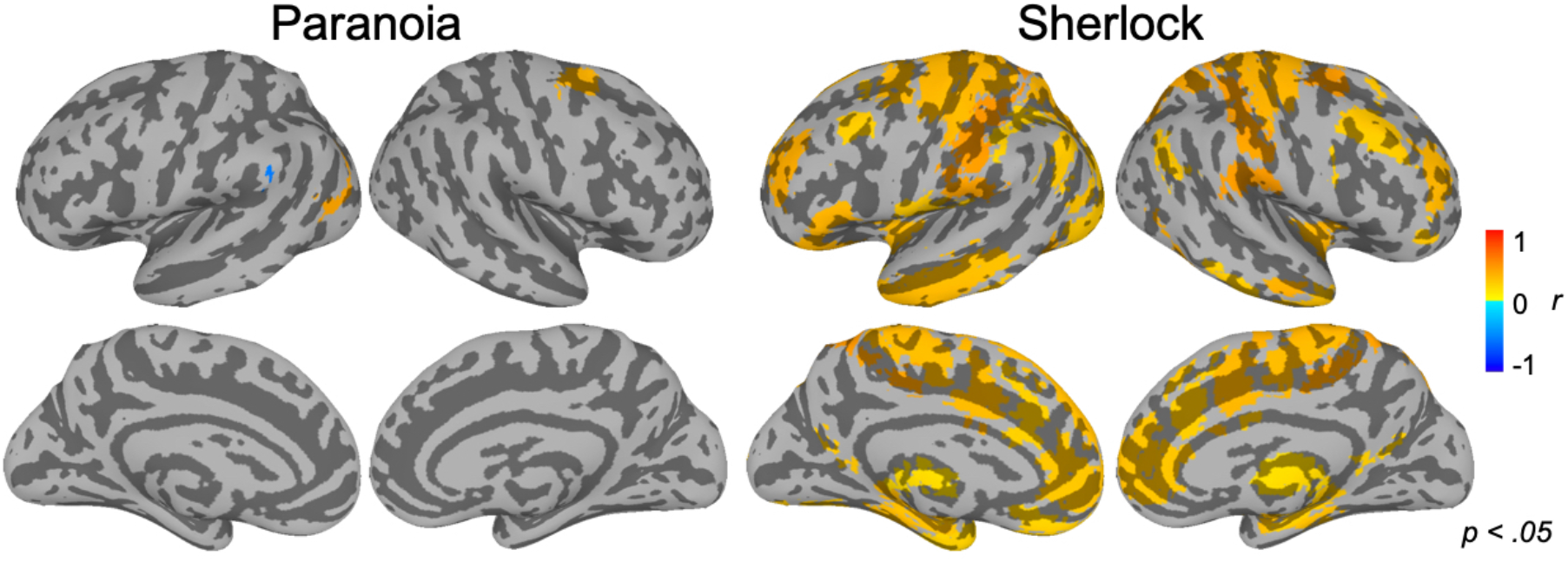
Regions that show significant correlations between dynamic ISC and sustained attention network strength, in Yeo atlas (two-tailed test; uncorrected *p* < .05).

**Supplementary Figure 6.**
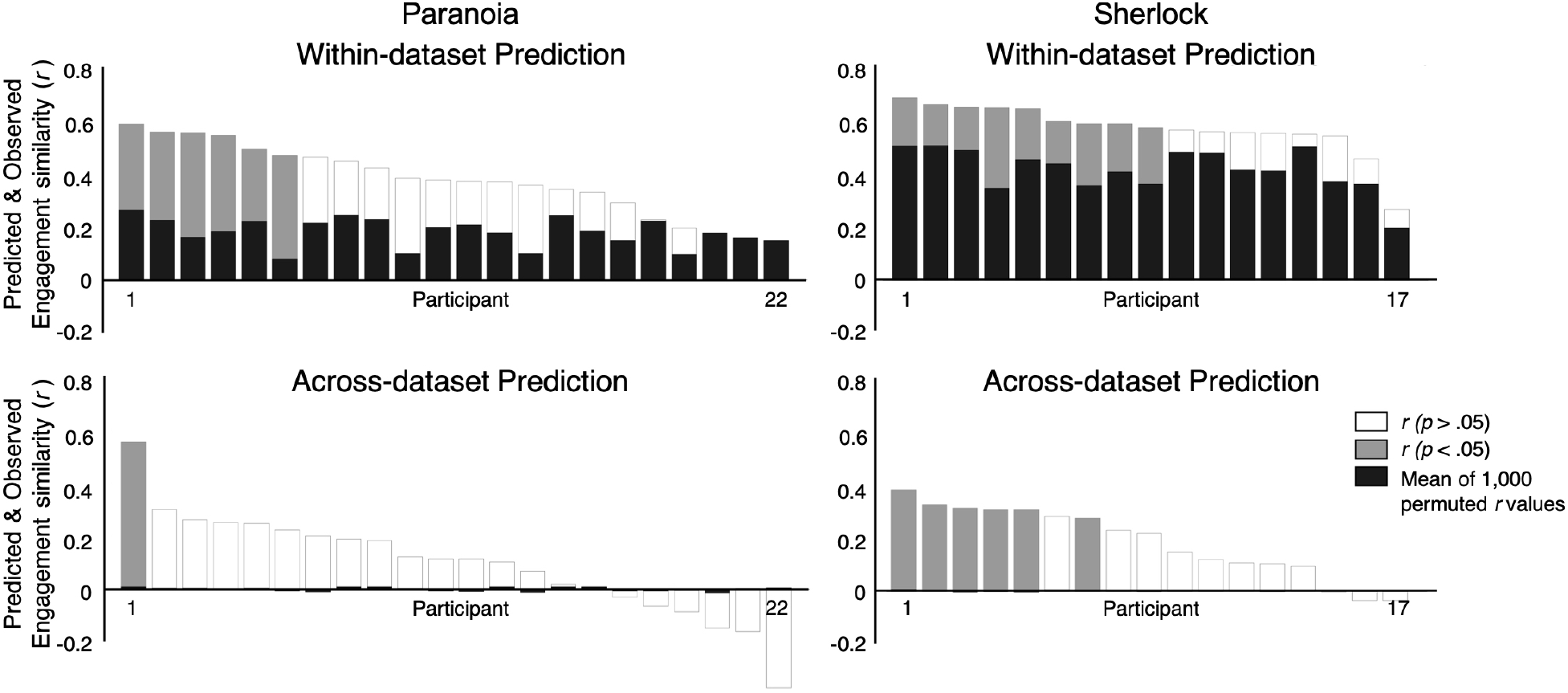
Prediction of group-average engagement time-course from patterns of functional connectivity. Prediction results for every cross-validation fold of the within-dataset (top) and across-dataset (bottom) predictions, respectively for *Paranoia* (left) and *Sherlock* (right). The black bars indicate the mean of the 1,000 permuted correlation values per fold. The gray bars indicate actual correlations, which were significantly larger than chance distributions (one-tailed test; uncorrected *p* < .05), and the white bars indicate actual correlations which did not pass significance threshold.

**Supplementary Figure 7.**
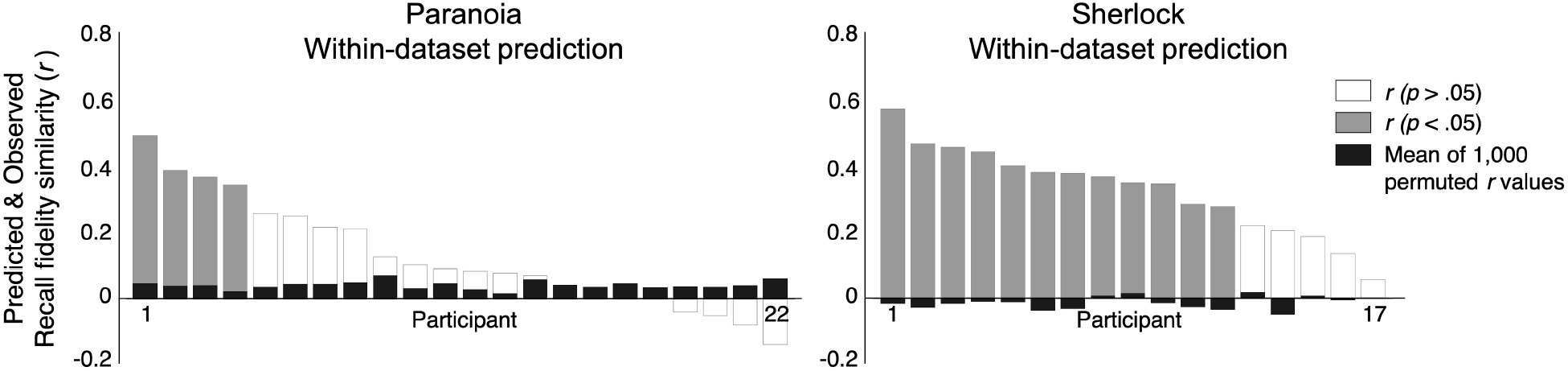
Prediction of individual-specific recall fidelity time-course from engagement network. Prediction results for every cross-validation fold of the within-dataset, respectively for *Paranoia* (left) and *Sherlock* (right). The black bars indicate the mean of the 1,000 permuted correlation values per fold. The gray bars indicate actual correlations, which were significantly larger than chance distributions (one-tailed test; uncorrected *p* < .05), and the white bars indicate actual correlations which did not pass significance threshold.

**Supplementary Table 1.**
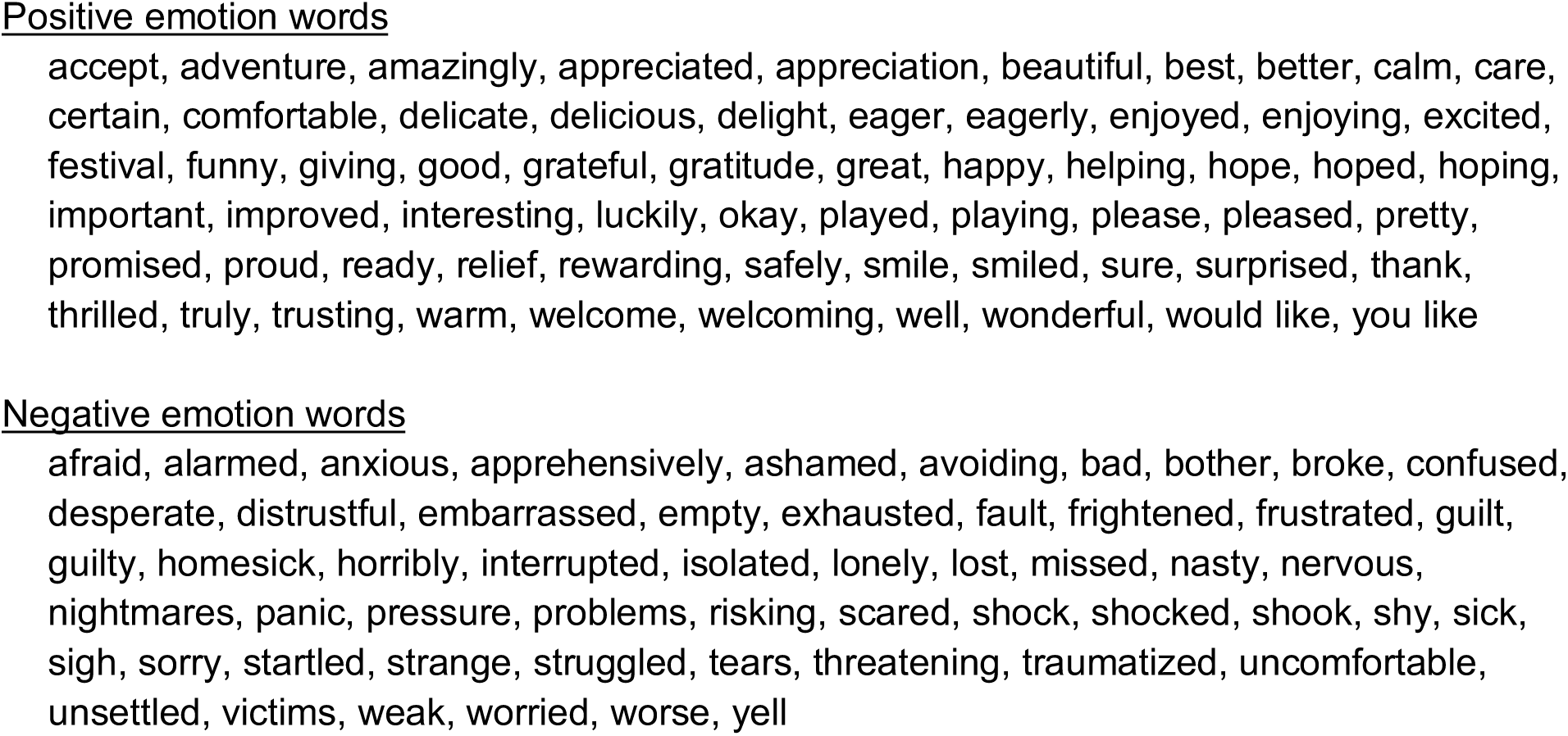
Lists of positively and negatively valenced words that appear in the *Paranoia* narrated transcript. We analyzed the Linguistic Inquiry and Word Count (LIWC; www.liwc.net) output of the *Paranoia* story transcript, provided by Finn et al.^18^ The LIWC software takes every word in the story transcript as input, and counts the number of words falling into different syntactic and semantic categories. The categories of our interests were ‘positive emotion’ and ‘negative emotion’.

**Supplementary Table 2.**
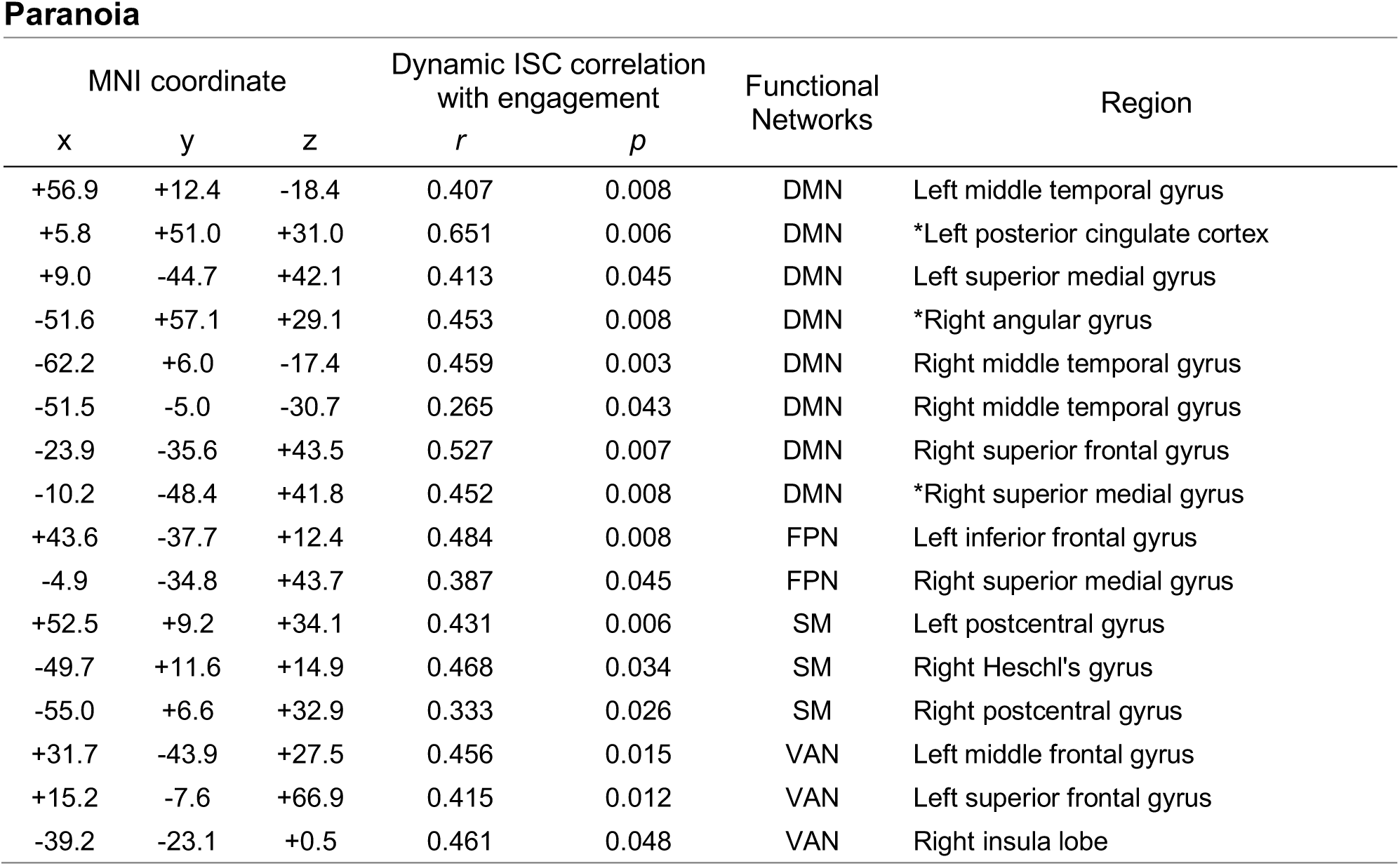
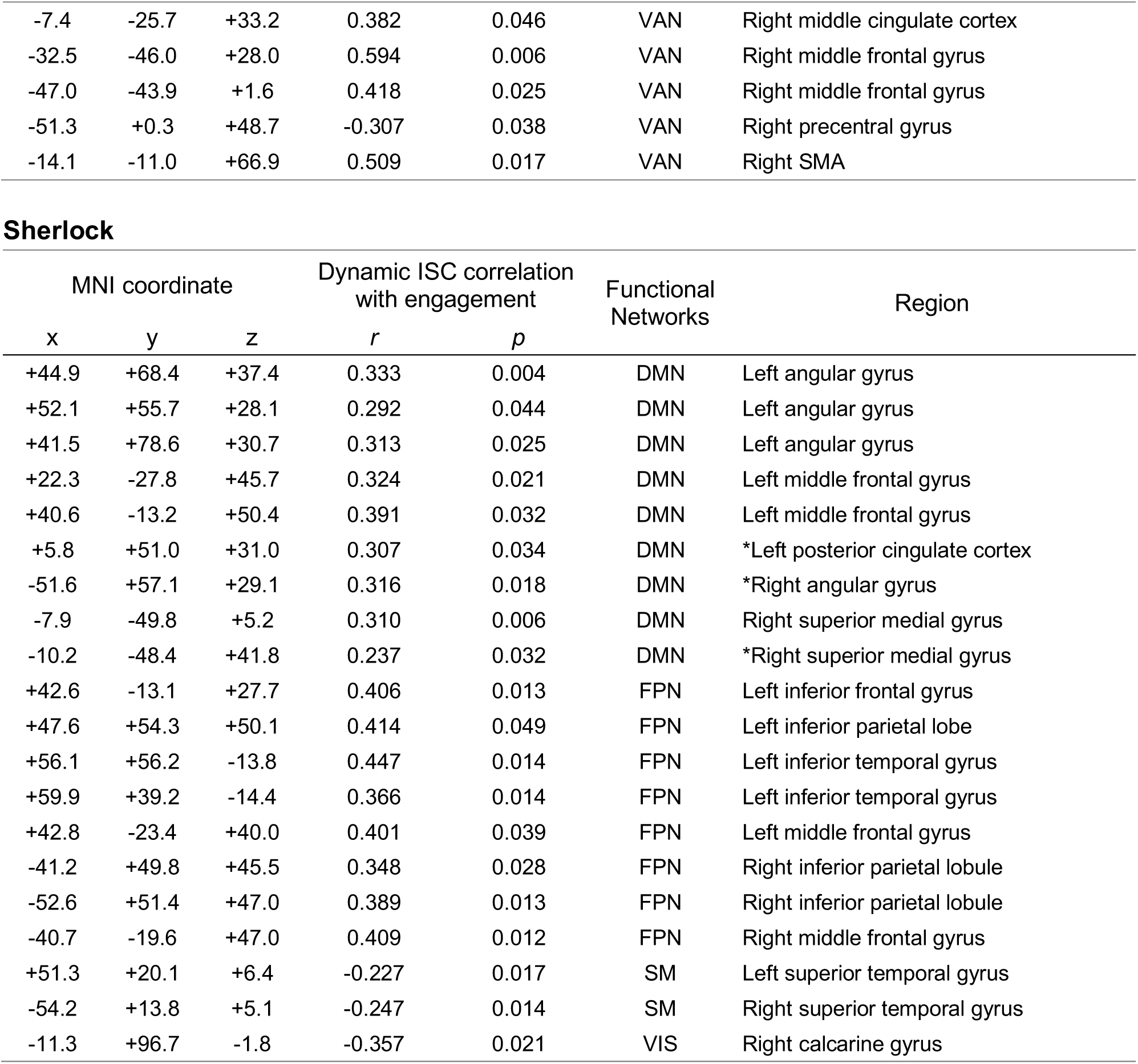
Regions that show significant correlation between dynamic inter-subject correlation (ISC) and group-average engagement, visualized in **Fig. 2b** (uncorrected *p* < 0.05), of *Paranoia* and *Sherlock* stories, respectively. The center of mass of each ROI was indicated by AFNI, and its coordinates and label were annotated within the MNI space, using Eickhoff-Zilles macro labels from N27. The *r* values indicate Pearson’s correlation of the ROI’s dynamic ISC and group-average engagement. The *p* values indicate an empirical *r*-value’s comparison with the null distribution where the engagement was phase randomized (two-tailed test, uncorrected for multiple comparisons). +Asterisks indicate regions that were significant in both datasets.

**Supplementary Table 3.** Anatomical overlap of the sustained attention network and the engagement network defined from *Sherlock* dataset, across different feature selection thresholds. In every round of leave-one-subject-out cross-validation, we applied feature selection to select relevant functional connections (FCs) of which time-courses are correlated with behavioral scores (i.e., gradCPT performance) or behavioral time series (i.e., group-average engagement). The FCs correlated with behavior—either in positive or negative direction—above significance threshold were selected. The FCs that were selected in every cross-validation was included in each network. We calculated the overlapping number of FCs showing the same directional correlations in the two networks. The main text uses threshold of *p* < .01. The significance of anatomical overlap was computed using the hypergeometric cumulative distribution function. (+): positively correlated with behavior, (−): negatively correlated with behavior.

| Feature selection threshold | Sustained attention network # of FCs | Engagement network # of FCs | # of overlapping FCs | Significance of overlap ($p$-value) |
| --- | --- | --- | --- | --- |
| .05 | (+) 321, (-) 505 | (+) 1218, (-) 193 | (+) 82, (-) 15 | (+) <.001, (-) .247 |
| .01 | (+) 134, (-) 200 | (+) 598, (-) 77 | (+) 20, (-) 0 | (+) .002, (-) .881 |
| .005 | (+) 95, (-) 134 | (+) 427, (-) 50 | (+) 11, (-) 0 | (+) .008, (-) .601 |
| .001 | (+) 45, (-) 56 | (+) 197, (-) 17 | (+) 3, (-) 0 | (+) .031, (-) .122 |

